# Display of the self-sufficient CYP102A1 on the surface of *E. coli*-derived Outer Membrane Vesicles

**DOI:** 10.1101/2021.06.02.446438

**Authors:** Delphine Devriese, Pieter Surmont, Frederic Lynen, Bart Devreese

## Abstract

The cytochrome P450 (CYP) monooxygenase superfamily offers the unique ability to catalyze regio-and stereospecifical oxidation of a non-activated C-H bond. CYPs found applications in the synthesis of pharmaceuticals and drug metabolites as well as in bioremediation. They are typically used in whole-cell bioconversion processes, due to their low stability and the need for a redox partner and cofactor. Unfortunately, substrate uptake and/or product transport limitations are frequently encountered and side reactions occur due to other enzymes in the cellular environment. Here, we present a proof-of-principle of a novel cell-free cytochrome P-450 nanocatalyst based on surface display on bacterial outer membrane vesicles. The self-sufficient CYP 102A1 from *Bacillus megaterium* was engineered to be translocated on the outer membrane vesicles of *Escherichia coli*. The resulting vesicles can simply be isolated from the culture supernatant. Moreover, no expensive and elaborate enzyme purification is required. This approach shows great promise as an alternative strategy to recombinantly produce CYP enzymes for a variety of applications, such as in fine chemical production and in the development of biosensors.

## 1 Introduction

Cytochrome P450 monooxygenases (CYPs) are versatile enzymes with a great potential in a variety of synthetic biology and biocatalyst applications [1]–[5]. CYPs contain a b-type heme group, linked to the enzyme via a coordinative bond between the heme iron and a conserved cysteine residue. The structure around this heme is highly conserved. This is in line with their common catalytic mechanism wherein they introduce one oxygen atom from molecular oxygen in a non-activated C-H bond while the other oxygen atom is reduced to water [6], [7]. Such a reaction is difficult to achieve by conventional chemical routes, typically requiring harsh conditions and rare metal catalysts. In contrast, CYPs are capable to perform these reactions in mild reaction conditions and often exhibit high regio-, chemo- and/or stereoselectivity, making them attractive biocatalysts [1]–[5]. Even though all CYPs share the same unique P450-fold consisting of a helix-rich α-region and a helix-poor β-region, some sites show high structural flexibility, enabling a high variability of substrates within this superfamily [6], [7]. Soluble CYPs from bacteria, as well as membrane-bound eukaryotic CYPs, are reported to catalyse industrially relevant reactions. A very successful CYP application is found in the production of artemisinic acid, the precursor of artemisin, an antimalarial drug. Fermentation titers of 25 g/l were obtained using an engineered *S. cerevisiae* strain, heterologously expressing *CYP71AV1* from the artemisin producing plant *Artemisia annua* [8].

Industrial application of CYPs is largely limited to whole-cell bioconversion approaches. Their use in enzyme reactors is limited due to their inherent instability and their requirement for redox partner(s) and the cofactor NAD(P)H. Using a whole-cell biocatalyst comes with some disadvantages as well. Interfering enzymes and pathways might be present, resulting in unwanted side reactions and/or by-products. The substrate and/or product may be toxic for the cells. Moreover, limitations regarding substrate uptake and/or product secretion occur. Several strategies have been investigated in order to overcome these disadvantages : strain engineering can be performed to knock out genes involved in unwanted side reactions, transporters can be introduced, cells can be permeabilized for increased substrate uptake, a two-phase system can be set up, etc. [1].

Engineering cells to translocate the enzyme to the extracellular side of the cell membrane combines some advantages of both fermentation and *in vitro* biocatalysis. By attaching the enzyme to the cell surface, substrate uptake and product transport limitations are circumvented, leading to higher productivities. Moreover, no elaborate and expensive enzyme purification is required. In the specific case of CYP production, the displayed enzyme is inserted *in vivo* in a membrane environment, enhancing stability. CYP102A1, the natural self-sufficient class VIII enzyme isolated from *Bacillus megaterium*, has been displayed on the surface of *E. coli* using three different approaches. In a first publication, Yim et al used the ice-nucleation protein (INP) InaK from *Pseudomonas syringae,* in a truncated form [9]. In a second publication, Ströhle et al reported the display of CYP102A1 by Autodisplay [10]. Thirdly, a recent article reports the display of CYP102A1 by *in situ* SpyCatcher-SpyTag interaction. [11].

Outer membrane vesicles (OMVs) offer a cell free alternative exploiting surface display. OMVs are spherical particles of 20-250 nm, released from the outer membrane (OM) by Gram-negative bacteria. Numerous functions have been ascribed to OMVs such as biofilm formation, pathogenesis and communication [12–14]. OMVs have emerged as a valuable vaccine delivery vehicle as they are able to modulate the immune response and much research has already been done to engineer these OMVs in order to obtain fit-for-purpose vesicles [15]. Next to vaccine delivery, OMVs have been suggested to serve as synthetic nanoreactors. Park et al were able to display a scaffold protein of 107 kDa on the surface of OMVs derived from *E. coli*, using an INP of *P. syringae*. Several enzymes were assembled on this scaffold, resulting in OMVs able to hydrolyse cellulose. The display on OMVs led to a marked activity increase when compared to the same scaffold, surface displayed on yeast cells [16]. It was hypothesized that this was due to the nanoscale dimension of the OMVs, leading to a higher enzyme:volume ratio and improved substrate accessibility, in contrast to the microscale dimension of a yeast cell [17].

In normal conditions, *E. coli* produces only a low amount of OMVs. For the use of OMVs as a biocatalyst, inducing a hypervesiculating phenotype is preferred [17]. OMVs are formed by blebbing of the OM. This blebbing is thought to occur at regions where peptidoglycan:membrane crosslinks are reduced, leading to decreased envelope stability [11], [18], [19]. Non-covalent interactions between the peptidoglycan (PG) layer and the Tol-Pal complex serves as one of the crosslinks, providing envelope stability. The Tol-Pal complex is composed of five proteins, that is TolA, TolB, TolQ, TolR and Pal, and spans the entire *E. coli* envelope. Deleting one or more of these Tol-Pal complex-encoding genes reduces the amount of crosslinks between the PG and the OM, consequently leading to increased vesiculation [20]. Next to the Tol-Pal complex, the deletion of other genes, encoding proteins involved in envelope crosslink modulation, were reported to result in a hypervesiculating strain, for example, knocking out the gene encoding the OM lipoprotein NlpI. It was hypothesized that this deletion resulted in a PG turnover imbalance. Proper crosslinking was therefore prevented, consequently leading to hypervesiculation [21], [22]. Furthermore, OMV biogenesis occurs as a response to external factors such as temperature, nutrient availability and quorum sensing [11], [18], [19]. Also, several antibiotics have been shown to induce OMV production. For example, subinhibitory concentrations of antimicrobial peptides which intercalate with the membrane, such as polymyxin B and colistin, lead to increased vesiculation, where the OMVs serve as a protective response [23].

We here explored whether the surface display of CYP102A1 could be taken one step further, i.e. if OMVs displaying CYP102A1 could be isolated, thereby creating a true *in vitro* cell-free biocatalyst. Several hypervesiculating mutants, as well as polymyxin B induced strains, were tested for their ability to produce OMVs, displaying this self-sufficient CYP and a proof-of-concept was delivered.

## 2 Material and methods

### 2.1 Materials

All chemicals were purchased from Sigma-Aldrich, unless specified otherwise.

### 2.2 Strains and media

For cloning as well as for surface display, the *E. coli* JM109 strain (New England Biolabs (NEB)) was used. Cells were grown in Luria-Bertani (LB) medium. LB consists of 1 % (w/v) NaCl (Merck), 0.5 % (w/v) yeast extract (Lab M) and 1 % (w/v) tryptone (Lab M). For agar plates, 1.5 % (w/v) agar (Lab M) was included. 100 µg/ml carbenicillin (Cb) (Gold Biotechnology) was added to medium and agar plates for selection of transformants and for plasmid maintenance during protein production.

For OMV display, several deletion mutants were used next to *E. coli* JM109 and are shown in Table 1. Two of the deletion mutants were ordered from the Keio collection [24]. In this collection, a duplicate of each deletion mutant is present. They include two independent mutants to account for handling errors, cross-contamination or accumulation of secondary mutations that. Both replicates for each deletion mutant were ordered for evaluation. For cloning, precultures and OMV quantification, cells were grown in LB medium. Strains from the Keio collection were grown in the presence of 25 µg/ml kanamycin (Kan) (Labconsult). In case of protein production, LB medium or Terrific Broth (TB) medium was used. TB consists of 1.2 % (w/v) tryptone (Lab M), 2.4 % (w/v) yeast extract (Lab M), 0.4 % (v/v) glycerol and phosphate buffer (0.017 M KH_2_PO_4_, 0.072 M K_2_HPO_4_). Alternatively, precultures were grown in non-inducing medium MDAG-135 and protein production was performed in autoinduction medium ZYM-5052, as described by Studier et al [25]. Here as well, 100 µg/ml Cb (Gold Biotechnology) was added to medium and agar plates for selection of transformants and for plasmid maintenance during protein production.

**Table 1:** *E. coli* deletion mutants, used for OMV display.

| Strain | Deletion | Source |
| --- | --- | --- |
| *tolA* 837 | $\Delta tolA$ | *E. coli* Keio Knockout Collection<br>Horizon Discovery: OEC4987-200826250 |
| *tolA* 838 | $\Delta tolA$ | *E. coli* Keio Knockout Collection<br>Horizon Discovery: OEC4987-213606638 |
| *nlpI* 863 | $\Delta nlpI$ | *E. coli* Keio Knockout Collection<br>Horizon Discovery: OEC4987-200828301 |
| *nlpI* 864 | $\Delta nlpI$ | *E. coli* Keio Knockout Collection<br>Horizon Discovery: OEC4987-213605299 |
| JC8031 | $\Delta tolRA$ | Described in [20]<br>Kindly provided by Prof. Dr. Laetitia Houot<br>(Laboratoire d'Ingénierie des Systèmes Macromoléculaires, CNRS) |

**Table 2:**
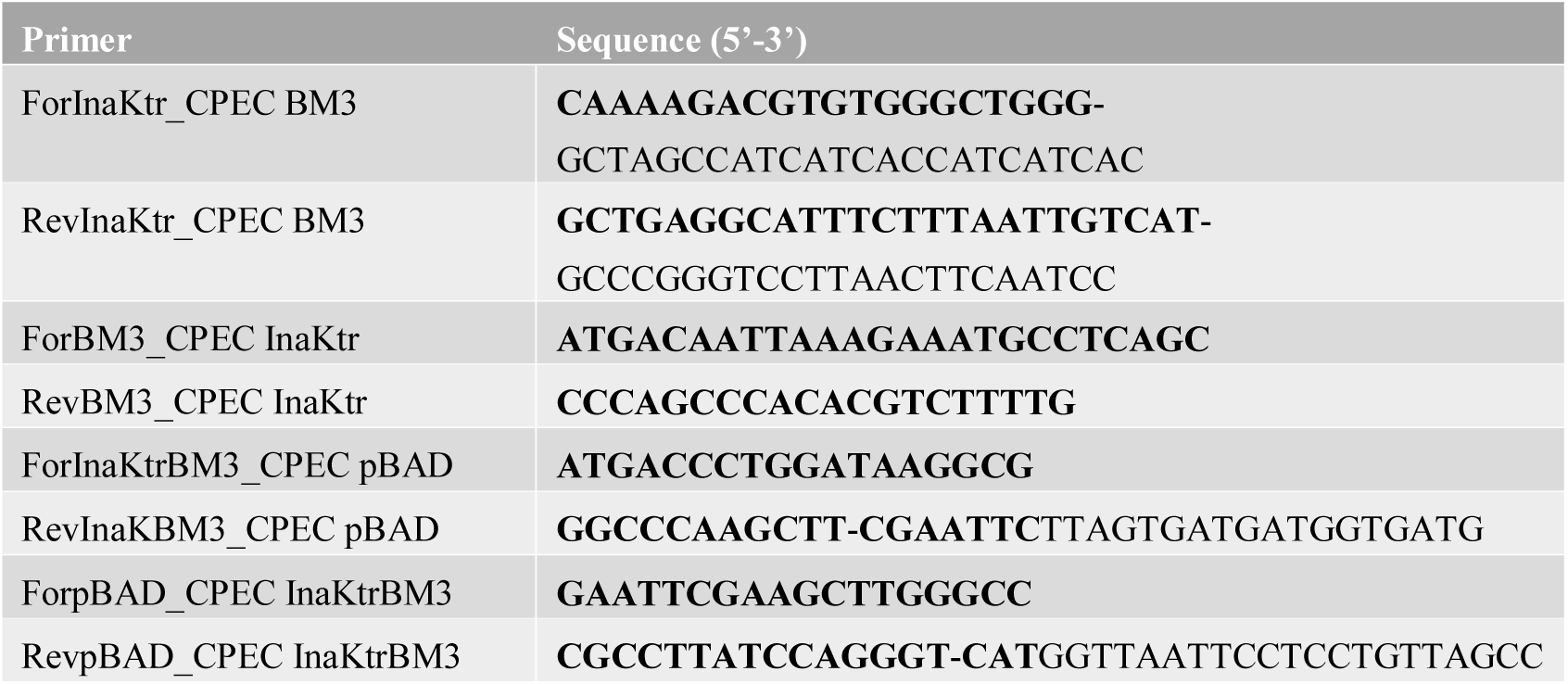
Primers used for molecular cloning of the different expression constructs, by means of CPEC. The bold sequence indicates the overlapping region between vector and insert. The hyphen separates the non-annealing part of the primer from the sequence annealing to the template for initial amplification of the respective inserts and linearization of the respective vectors.

### 2.3 DNA constructs and molecular cloning

InaK was truncated for its use as a surface display anchor, and is from here on now denoted as InaKtr. A construct was designed, consisting of the N- and C-terminal unique domains and a C-terminal Asp-Pro-Gly linker, as described by Yim et al [9]. Truncation of the central domain was performed based on the previous report of Shimazu et al [26], who deleted the central domain except for subdomain 1 (an incomplete 32-residue repeat, which may serve as a transition between the N-terminal and the repeating domain) and one 48 residue repeat from subdomain 4. Furthermore, a NheI restriction site and sequence encoding a Histag of six His residues at the C-terminus were introduced. This construct was codon optimized, synthesized and cloned in the vector pMAL-c4x by GenScript. Cloning was performed in between restriction sites NdeI and EcoRI, thereby excluding the maltose binding protein encoding gene. The pMAL-c4x vector was chosen for its *tac* promotor, as this promotor was proposed to be optimal for recombinant CYP expression in *E. coli* [27], [28]. The construct will further be referred to as InaKtr_pMAL (Fig. S1).

For all molecular cloning experiments, Circular Polymerase Extension Cloning (CPEC), was applied. All primers are given in **Error! Reference source not found.**. The gene encoding CYP102A1, otherwise denoted as BM3, was amplified from genomic DNA of *Bacillus megaterium* (kindly provided by Prof. Inge Van Bogaert (CSB, Ghent University)), using the primer pair ForBM3_CPEC InaKtr - RevBM3_CPECInaKtr. The InaKtr_pMAL vector was linearized and amplified using the primers ForInaKtr_CPECBM3 and RevInaKtr_CPECBM3. The amplified *bm3* gene was subsequently inserted in this linearized InaKtr_pMAL vector, in frame of the sequence encoding the Asp-Pro-Gly linker, preceding the NheI restriction site and the Histag encoding sequence, resulting in BM3 with an N-terminal InaKtr surface display anchor and a C-terminal Histag. The construct will be denoted as InaKtrBM3_pMAL.

In order to place the expression of *inaKtrBM3* under the control of the *araBAD* promotor, the complete coding sequence was amplified with the primers ForInaKtrBM3_CPECpBAD and RevInaKtrBM3_CPECpBAD (**Error! Reference source not found.**) for insertion in the vector pBAD/Myc-HisA. The vector was amplified and linearized using the primers ForpBAD_CPECInaKtrBM3 and RevpBAD_CPECInaKtrBM3 (**Error! Reference source not found.**).

### 2.4 Culture conditions

All cultures were started with a 5 ml preculture using LB medium including the appropriate antibiotics in a 50 ml shake flask, which were incubated overnight at 37 °C and 200 rpm. In case of the OMV experiments, 1 % glucose was added for the pMAL constructs.

In case of surface display, *E. coli* JM109 was grown in 50 ml LB including 0.5 mM δ-aminolevulinic acid (5-ALA) and 100 µg/ml Cb, inoculated with 500 µl preculture, in a 500 ml shake flask at 28 °C and 200 rpm until an OD_600_ of 0.5. At this point, expression was induced by the addition of isopropyl-β-D-thiogalactoside (IPTG) (Gold Biotechnology) to a final concentration of 1 mM. The expression phase was carried out overnight and cells were collected the next day for subsequent assays and analyses.

For OMV quantification, the respective *E. coli* strains were grown in 50 ml LB (including Kan in case of strains from the Keio collection), in a 500 ml shake flask after inoculation with 500 µl preculture. The cultures were incubated at 37 °C and 200 rpm until an OD_600_ of 0.5, at which point the supernatant was collected for sOMV isolation as described below. Alternatively, cultures were grown overnight for sOMV collection in the stationary phase. In case of quantifying the polymyxin B induced hypervesiculation, JM109 was grown in 100 ml LB in a 1 l shake flask at 37 °C and 200 rpm until an OD_600_ of 0.5. The culture was split in two and polymyxin B (Labconsult) was added to a final concentration of 0.75 µg/ml. Incubation continued for another 3 h and sOMVs were collected.

When OMV display was pursued, the respective *E. coli* strains were grown in 100 ml LB or TB with 0.5 mM 5-ALA and 100 µg/ml Cb in a 1 l shake flask. For strains from the Keio collection, Kan was added and for the pMAL constructs, 0.5 % glucose was included.

Cultures were incubated at 37 °C and 200 rpm until an OD_600_ of 0.5. At this point, the temperature was lowered to 28 °C and expression was induced either by IPTG addition to a final concentration of 0.3 mM (pMAL constructs) or by arabinose addition to a final concentration of 0.2 % (w/v) (pBAD constructs). Expression was carried out to the late exponential phase. In case of expression in *E. coli nlpI* 863, an overnight expression phase was performed. The cells and/or supernatant were subsequently collected for OMV isolation. In early experiments, expression was induced with 1 mM IPTG, as indicated in the results section. No glucose was added in these cultures. Other expression times have been used, as indicated in the results section. In those experiments, the growth phase was carried out at 28 °C as well.

### 2.5 Cell lysis and outer membrane collection

Cells were collected by centrifugation at 20 000 g and 4 °C for 15 min. The supernatant was discarded and the cell pellet was resuspended in 50 mM Tris-HCl, pH 7.4, including protease inhibitor. Cell lysis was done by sonication on ice. Following lysis, the cell debris and unlysed cells were pelleted by centrifugation. A second sonication step was included and the two supernatant lysate samples were pooled for membrane collection by ultracentrifugation at 100 000 g and 4 °C for 1 h. The membrane pellet was washed with the same buffer, including 300 mM NaCl (Merck), and ultracentrifugation was repeated. Collection of the outer membrane (OM) was performed as described by Sandrini et al [29], with modification. To solubilize the inner membrane (IM), the washed membrane pellet was incubated with 50 mM Tris-HCl, pH 7.4, including 2 % Triton X-100 for 30 min at room temperature with occasional mixing. A last ultracentrifugation step at 100 000 g and 4 °C for 1 h was performed, resulting in an OM pellet and solubilized IM in the supernatant.

### 2.6 OMV collection

#### 2.6.1 sOMV collection

Spontaneously formed outer membrane vesicles (sOMVs) were isolated from the culture supernatant after pelleting the cells by centrifugation at 10 000 g and 4 °C for 10 min. The collected supernatant was filtered using a pore size of 0.22 µm. In later experiments, a switch was made to a 0.45 µm filter, as indicated in the results section. Next, the filtered medium was ultracentrifuged for 3 h at 100 000 g and 4 °C. The OMV pellet was washed with phosphate buffered saline (PBS), followed by ultracentrifugation. The final OMV pellet was resuspended in PBS for subsequent enzymatic assays and/or analyses. For OMV quantification, no washing step was included and the obtained pellet was immediately resuspended in 1 ml PBS.

#### 2.6.2 eOMV extraction

For EDTA extraction of outer membrane vesicles (eOMVs), the method described by Van de Waterbeemd et al [30] was used, with modifications. Cells were collected by centrifugation at 10 000 g and 4 °C for 10 min. The cell pellet was resuspended in 7.5 volumes (ml/g wet weight) 100 mM Tris-HCl, pH 8.6, 10 mM EDTA, and incubated for 30 min at room temperature. Cells were pelleted by centrifugation and the supernatant was collected. Subsequently, filtration was performed applying a pore size of 0.22 µm. Alternatively, a pore size of 0.45 µm was chosen, as indicated in the results section. eOMVs from the supernatant were pelleted by ultracentrifugation for 3 h at 100 000 g and 4 °C. Similar as for the sOMVs, wash and final resuspension was performed in PBS.

### 2.7 SDS-PAGE and western blot

Protein samples were mixed with Laemmli buffer and the proteins were separated by SDS-PAGE using either a 7.5 % or a 12 % polyacrylamide gel and a Tris-glycine running buffer (Bio-Rad). The Precision Plus Protein™ Unstained Standard (Bio-Rad) was used as a molecular weight marker. Gels were stained with Coomassie brilliant blue G.

In case of western blot analysis, SDS-PAGE was performed as described above, using the Precision Plus Protein™ Dual Color Standard (Bio-Rad). After protein separation was completed, the proteins were transferred to a nitrocellulose blotting membrane. The membranes were incubated with a 5 % (w/v) nonfat dry milk (Bio-Rad) solution in PBS and washed with PBS-T (PBS including 0.1 % (v/v) Tween-20) before incubation with a horseradish peroxidase (HRP)-conjugated 6x-His epitope tag monoclonal antibody, diluted in a 1 % (w/v) nonfat dry milk (Bio-Rad) solution in PBS-T. The anti-His(C-term)-HRP antibody (R931-25, Invitrogen) was used, at a dilution of 1:10 000. After washing with PBS-T, proteins were visualized by chemiluminescence.

### 2.8 Trypsin digestion and MALDI-TOF MS

Gel bands of interest were cut out after separation by SDS-PAGE and visualization of the proteins with Coomassie brilliant blue G. After complete removal of the Coomassie stain using 50 % acetonitrile (ACN) (Biosolve) in 200 mM NH_4_HCO_3_ at 30 °C, a reduction step with 10 mM DTT in 100 mM NH_4_HCO_3_ for 1 h at 56 °C and an alkylation step with 55 mM iodoacetamide (IAA) in 100 mM NH_4_HCO_3_ for 45 min at room temperature in the dark was performed. Gel bands were subsequently washed with 100 mM NH_4_HCO_3_ and dehydrated with 100 % ACN (Biosolve) (two times). Following complete dehydration of the gel bands, 0.02 µg modified trypsin (Promega) in 10 µl 50 mM NH_4_HCO_3_ was added and samples were incubated on ice for 45 min. 40 µl 50 mM NH_4_HCO_3_ was added and the proteins were digested overnight at 37 °C. The next day, peptides were extracted twice with a 60 % ACN (Biosolve) solution containing 0.1 % formic acid. The extracts were pooled and dried under vacuum.

The dried peptides were then resuspended in 10 µl of a 50 % ACN (BioSolve) solution containing 0.1 % formic acid. 1 µl of resuspended peptides, mixed with a saturated α-cyano-4-hydroxycinnamic acid solution in a 1:1 ratio was spotted onto an Opti-TOF 384 Well MALDI Plate Insert for Matrix-Assisted Laser Desorption Ionization Time-of-Flight Mass Spectrometry (MALDI-TOF MS) analysis with the MALDI TOF/TOF 4800 Plus (ABSciex). Identification was done with Mascot using the *E. coli* (strain K12) protein database, downloaded from Uniprot.

### 2.9 Fatty acid bioconversion assay

Hydroxylation of fatty acids (FAs) was assayed as follows. The reaction was performed in a volume of 1 ml, containing 100 mM potassium phosphate, pH 7.4, 0.1 mM palmitic acid (from a 10 mM stock in DMSO), 1 U glucose-6-phosphate dehydrogenase (G6PDH), 5 mM glucose-6-phosphate (G6P) and 1 mM NADPH. The reaction was carried out for 20 min at 37 °C and 500 rpm. The conversion was subsequently quenched with HCl and extraction was performed with and equal volume of diethyl ether (three times). The diethyl ether extracts were pooled, evaporated under vacuum and stored at – 20 °C until GC-MS analysis.

### 2.10 GC-MS analysis

The evaporated extract was dissolved in 50 µl 1 % trimethylchlorosilane (TMCS) in N,O- bis(trimethylsilyl) trifluoroacetamide (BSTFA) and incubated for 30 min at 75 °C. After a centrifugation step at 10 000 g for 10 min, the supernatant was transferred to a glass vial for GC-MS analysis. The GC-MS system used was the 7890-5975C from Agilent in combination with the HP-5MS column (5% phenyl methyl polysiloxane, 30m×0.25mm ID, 0.25µm) from Agilent. The temperature was maintained at 180 °C for 1 min, subsequently raised to 300 °C at 8 °C/min and then held isotherm for 5 min.

### 2.11 12-pNCA assay

After surface display, the cells were collected by centrifugation at 10 000 g and 4 °C for 10 min and washed three times with PBS. Thereafter, the cells displaying InaKtrBM3 were resuspended in PBS to a final OD_600_ of 50. OMVs were assayed after collection as described above. For measuring the activity of InaKtrBM3, the assay developed by Schwaneberg et al [31] was applied, with modifications. The reaction was carried out in a total volume of 250 µl, consisting of 100 mM Tris-HCl, pH 8.2 and either cells at a final OD_600_ of 2 or 25 µl resuspended OMVs. To this reaction, *p*-nitrophenoxydodecanoic acid (12-pNCA) (Molport) was added to a final concentration of 50 µM from a 5 mM stock solution in DMSO. After preheating at 30 °C, NADPH was added simultaneously to a final concentration of 0.1 mM. The formation of *p*-nitrophenol was monitored at 405 nm (ε = 13 200 M^-1^cm^-1^) and 30 °C for 30 min with shaking with the Bio-Rad Microplate Reader model 680.

### 2.12 Nanoparticle Tracking Analysis

OMVs were quantified by means of nanoparticle tracking analysis (NTA), using the NanoSight LM10-HS microscope (Malvern Instruments Ltd), equipped with a 405 nm laser. The OMV suspension was diluted with PBS to fit within the measuring range of 3 x 10^8^ and 1 x 10^9^ particles/ml. For every sample, three movies of 30 s were recorded and the temperature was monitored. The recorded movies were processed with the NTA Analytical Software version 2.3. Three biological replicates were measured for assessing hypervesiculation. Subsequent statistical analysis was performed using the Graphpad Prism software.

## 3 Results

### 3.1 The surface display construct InaKtrBM3 is active

We initiated the development of OMVs displaying CYP102A1 (also denoted as BM3) by the construction of the vector, containing the InaKtrBM3-encoding sequence for the surface display of BM3. Subsequently, a whole-cell bioconversion assay was performed, assaying the hydroxylation of palmitic acid by the intact cells. Following the bioconversion assay, the products were extracted and silylated for GC-MS analysis. The chromatogram obtained after a 20 min palmitic acid conversion by surface displayed InaKtrBM3 is shown in Figure.1. Two small peaks appeared in this chromatogram, i.e. at RT 7.971 min and 8.105 min. The MS spectra of both peaks are shown in Figure 2. Both spectra coincide with hydroxylated palmitic acid. The molecular ion of m/z 416 is not found, and it indeed is reported that the molecular ion of silylated OHFAs is only present in low abundance [32]. Instead the ion [M-15]^+^, missing one methyl group, and the ions [M-31]^+^ and [M-105]^+^, resulting from interactions between the two functional groups, are seen. In the MS spectrum of Figure 2A, the ion with m/z 131 is highly abundant, indicating that palmitic acid is hydroxylated at the ω-2 position. The spectrum of Figure 2B shows a highly abundant ion with m/z 117, indicating hydroxylation at the ω-1 position. InaKtrBM3 thus actively converts palmitic acid mainly to 14-, and 15-hydroxypalmitic acid. This experiment showed that CYP102A1 was successfully displayed at the cell surface of *E. coli* in an active form.

**Figure.1:**
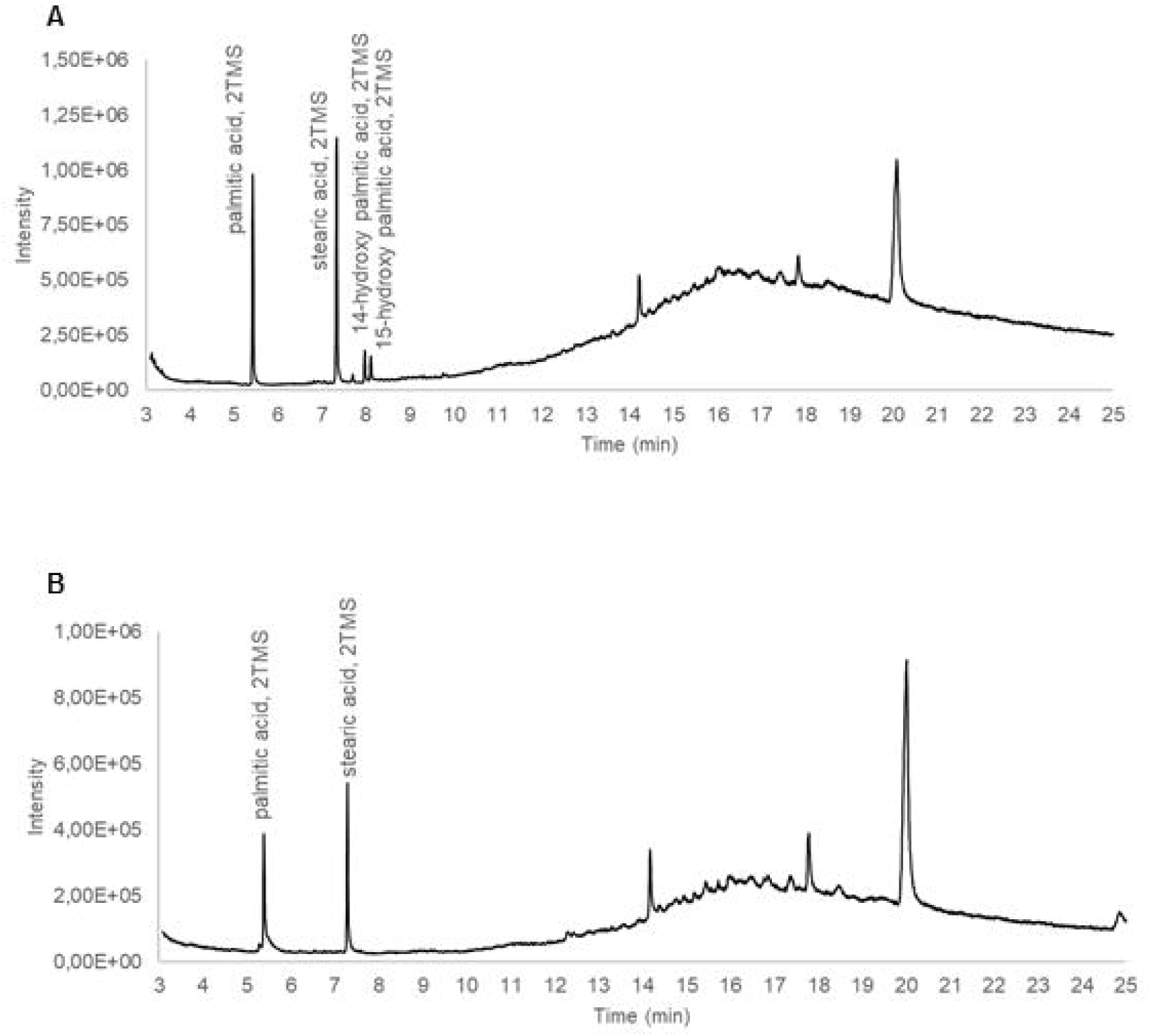
A: GC chromatogram of silylated extracts after a 20 min whole-cell palmitic acid bioconversion assay using JM109 cells, surface displaying InaKtrBM3. Palmitic acid was hydroxylated at its ω-1 and ω-2 position. B: GC chromatogram of silylated extracts of a blank, i.e. no palmitic acid substrate was added during bioconversion. The identified palmitic acid and stearic acid are contaminants, always observed by GC-MS analysis. Only peaks identified as silylated compounds, based on the presence in the MS spectrum of the characteristic peaks at m/z 73 and 75, are annotated.

**Figure 2:**
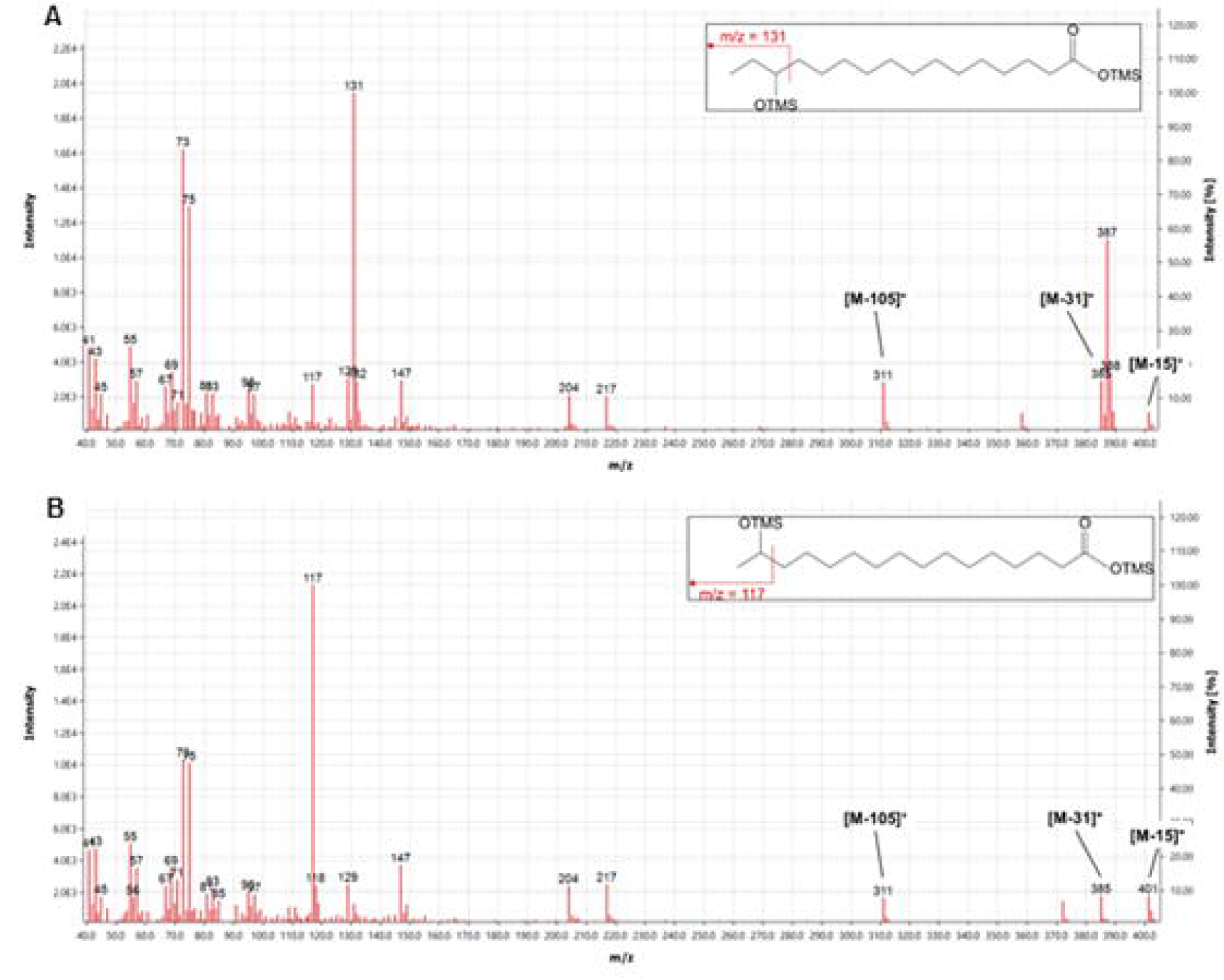
MS spectra of hydroxylated palmitic acid. A: MS spectrum of peak at RT 7.971 min (Figure.1), that is 14-hydroxypalmitic acid. B: MS spectrum of peak at RT 8.105 min (Figure.1), that is 15-hydroxypalmitic acid.

### 3.2 Evaluation of hypervesiculating *E. coli*

Interest went to creating OMVs, displaying the self-sufficient CYP, in order to obtain a true *in vitro* system, with these OMVs serving as a nanoscale carrier. In normal conditions, *E. coli* produces only a low amount of OMVs. It is thus desired to induce hypervesiculation for an increased OMV yield [17]. Several mutations, as well as membrane intercalating molecules, have been reported to result in increased vesiculation. Single deletion mutants were ordered from the Keio collection and it was evaluated whether these indeed led to increased OMV formation. The single deletion mutant *tolA* was selected for the first OMV display experiments. Additional strains were ordered, i.e. the two independent *nlpI* deletion mutants. Of special interest was the hypervesiculating *E. coli* strain JC8031, where both *tolA* and *tolR* are knocked out. This strain was reported to show a marked vesiculation increase. Additionally, this strain proved to be successful in the OMV display of recombinant proteins in several studies [16], [33], [34]. OMVs were isolated when the cultures were in the early exponential phase and quantified by NTA (**Error! Reference source not found.**A). In the early exponential phase, both *tolA* 838 and JC8031 produce a significantly higher amount of OMVs. Neither of the *nlpI* deletion mutants show hypervesiculation, this in contrast to previous reports [21], [22]. However, in these studies, OMVs were quantified after collection from a stationary phase culture. Schwechheimer et al hypothesized that NlpI suppresses PG endopeptidase activity. Deletion of *nlpI* leads to uncontrolled PG degradation, to which the cell responds by an increased PG synthesis, even in the stationary phase, in order to survive. In the same study, it was observed that NlpI was specific for the PG endopeptidase Spr in the stationary phase, whereas the penicillin binding protein PBP4 was supressed in the exponential phase. Due to these differences between the exponential and stationary phase, OMVs from the different deletion mutants were isolated and quantified after overnight growth to the stationary phase (**Error! Reference source not found.**B). Indeed, both *nlpI* deletion mutants now showed to produce an increased amount of OMVs, compared to JM109.

Several antibiotics have been shown to induce OMV production. It was investigated whether the JM109 strain showed increased OMV production adding polymyxin B. The subinhibitory concentration of 0.75 µg/ml, as applied by Manning and Kuehn [23], was added to an exponential phase culture of JM109. After an incubation time of 3 h, the medium was collected for OMV isolation and quantification by NTA (Figure 4). Indeed no growth inhibition was observed using this polymyxin B concentration, as the OD_600_ did not differ after a 3 h incubation (Figure 4A). Growth inhibition was observed only from a concentration of 1.5 µg/ml on (data not shown). Upon incubation with polymyxin B, a significantly increased amount of OMVs was observed (Figure 4B). However, when compared to spontaneous release of OMV by JC8031, this increased vesiculation appeared to be negligible (Figure 4C). This confirms previous reports that factors such as antibiotics and PQS do not increase the vesiculation to yields, useful for pharmaceutical and biotechnological applications [17].

**Figure 3:**
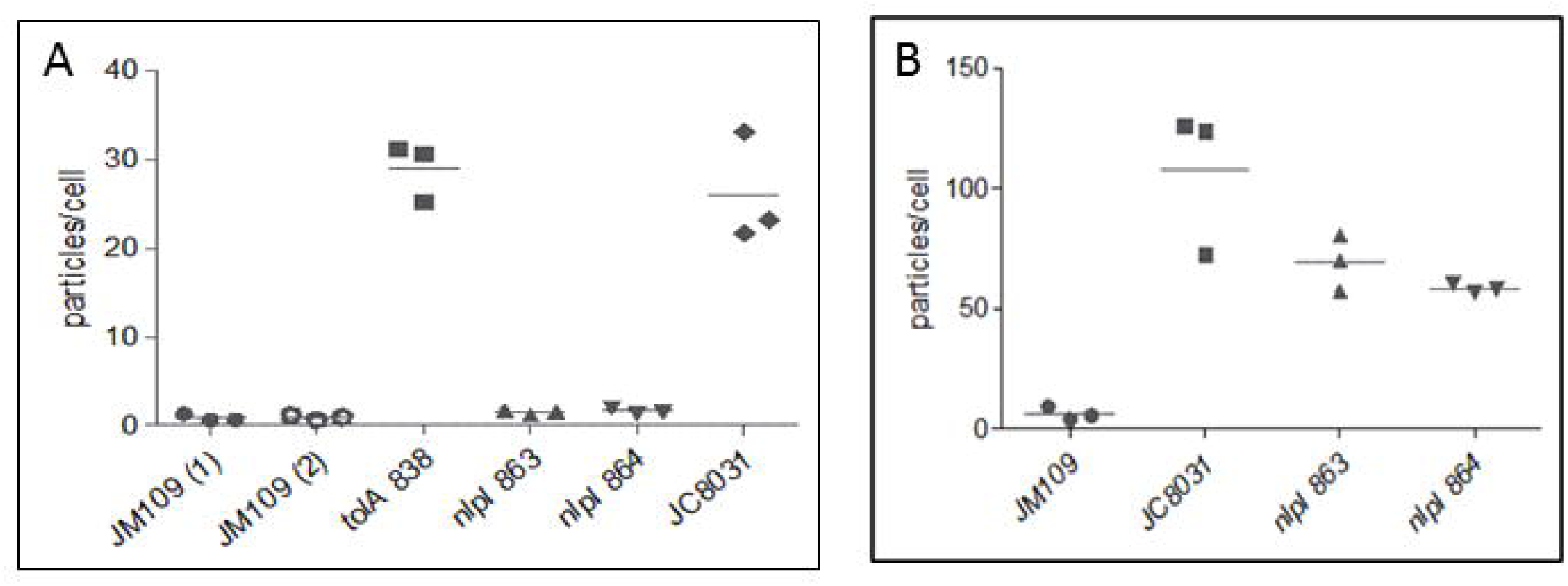
OMV quantification by NTA, displayed as particles per cell. A: OMVs were collected in the exponential phase. Three biological replicates were taken along in parallel. The vesiculation was significantly different between the different strains, as determined by one-way ANOVA (p < 0.0001). Tukey’s multiple comparison test showed that both the *tolA* deletion mutant and JC8031 produced a significant higher amount of vesicles compared to all other strains, but did not differ significantly from each other. The three biological replicates at timepoint 1, that is JM109 (1), did not differ from the three biological replicates at timepoint 2, that is JM109 (2), proving reproducibility. B: OMVs were collected after overnight growth, that is in the stationary phase. Three biological replicates were taken along in parallel. The vesiculation was significantly different between the different strains, as determined by one-way ANOVA (p = 0.0004). All three deletion mutants showed a significantly higher vesicle production compared to JM109, based on Tukey’s multiple comparison test. JC8031 showed to vesiculate to a higher extend than *nlpI* 864, but not *nlpI* 863, based on Tukey’s multiple comparison test. *nlpI* 863 and 864 vesiculation did not significantly vary.

**Figure 4:**
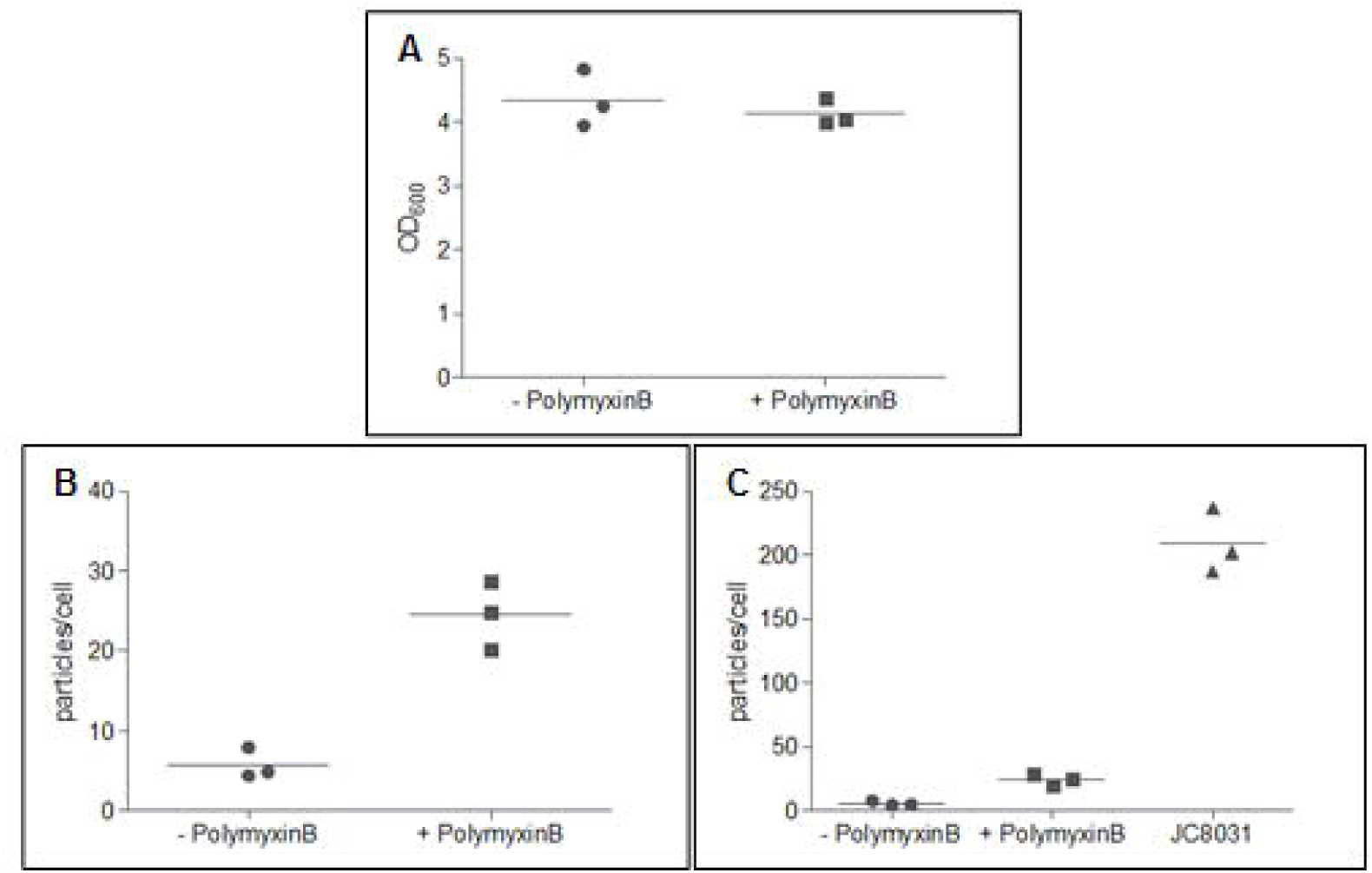
Polymyxin B induced vesiculation. A: Optical density of JM109 after 3 h growth post polymyxin B addition to one of two cultures, executed in triplicate. The optical density did not differ significantly in absence or in presence of 0.75 µg/ml polymyxin B, determined by an unpaired t-test (p = 0.5262). B: OMV quantification by NTA, displayed as particles per cell, after induction of OMV biogenesis by polymyxin B addition, compared to a non-induced control. Polymyxin B induction significantly increased vesiculation, determined by an unpaired t-test (p = 0.0022). C: OMV quantification by NTA, displayed as particles per cell. OMVs were collected after growing the JC8031 strain for the same period of time as the polymyxin B induced cultures (performed in triplicate). The vesiculation increase by polymyxin B addition became insignificant when including the hypervesiculating strain, determined by one-way ANOVA (p < 0.0001), followed by Tukey’s multiple comparison test.

### 3.3 The *tolA* deletion mutant did not result in OMVs displaying InaKtrBM3

Initial experiments were performed with the *tolA* deletion mutants (i.e. *tolA 837* and *tolA 838*). A first 50 ml expression test was performed where *tolA* 837 and 838 were grown in parallel. InaKtrBM3 production was induced at an OD_600_ of 0.5 with 1 mM IPTG and the cells were grown further to the late exponential phase, at which point the OMVs were isolated and analysed by western blot (Figure 5A). No signal was observed, so no OMVs displaying InaKtrBM3 were isolated. Continuing with *tolA* 838, some alternative expression conditions were investigated. Additionally, in all conditions, a larger culture was grown in order to increase the amount of isolated OMVs. In a first setup, OMVs were isolated after an expression phase of 3 h. Secondly, an overnight expression phase was performed. Thirdly, instead of IPTG induction, cells were grown in autoinduction medium, as it has been stated to lead to increased densities [25], [35]. Lastly, overnight expression at 20 °C instead of 28 °C was performed in order to assess whether this would lead to improved yield of properly folded enzyme and thus increase the yield displayed on OMVs. Unfortunately, none of these conditions led to successful isolation of OMVs displaying InaKtrBM3, as no signal was observed by western blot analysis (Figure 5B). To verify whether the enzyme was effectively produced and surface displayed, the OM was isolated and analysed in parallel. In all cases where expression was induced with IPTG, InaKtrBM3 was present around the right MW (152 kDa), together with several degradation bands. These degradation bands could partially be ascribed to proteolysis of the linker sequences. Proteolysis of the linker sequence in between InaKtr and BM3 leads to a signal at 118 kDa. Release of the BMR sequence by proteolysis of the linker in between the heme and reductase domain theoretically gives a signal around 65 kDa. In case of autoinduction, only one band is seen and this at a lower MW. Autoinduction thus appears to be unsuitable for surface display of this enzyme.

**Figure 5:**
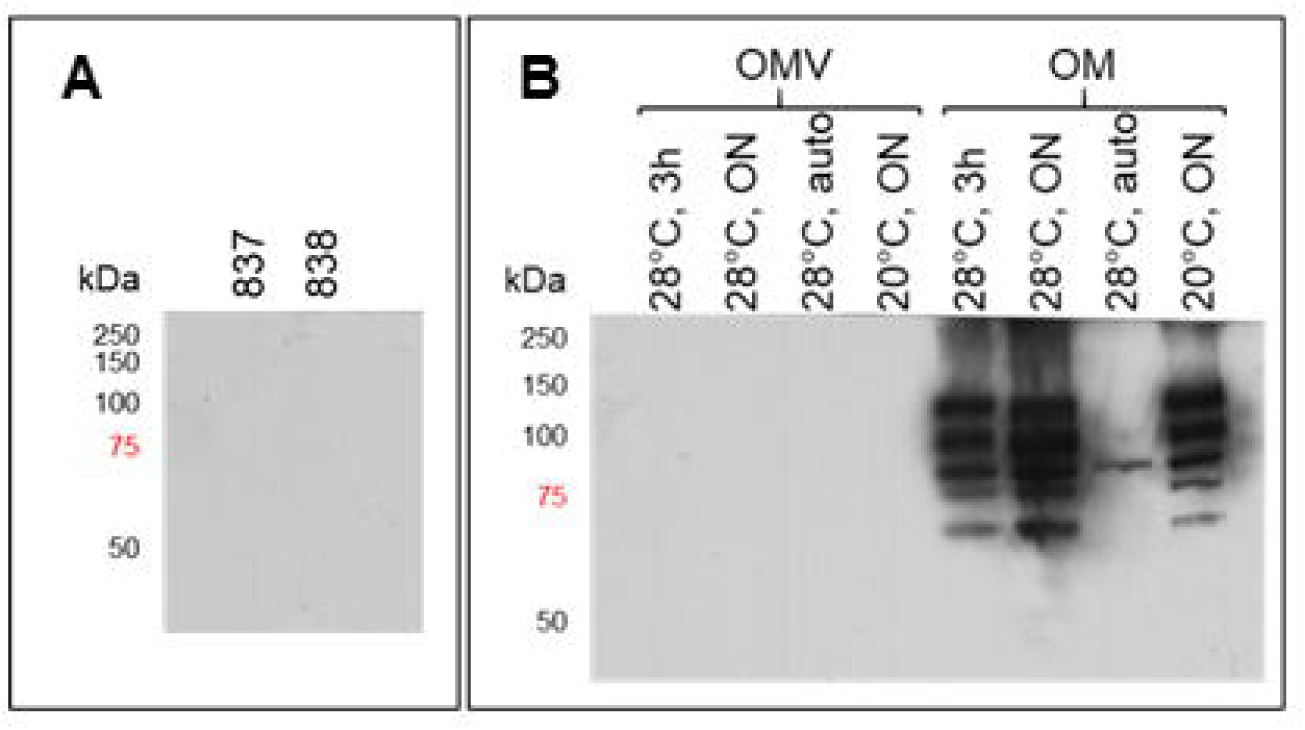
Western blot analysis after InaKtrBM3 (152 kDa) production in the *tolA* deletion mutant(s). A: OMV samples isolated after expression in both *tolA* 837 and *tolA* 838. B: OMV and OM samples after InaKtrBM3 production in *tolA* 838, applying four different expression conditions.

### 3.4 JC8031 did not result in OMVs displaying InaKtrBM3 at first

No success was booked using the *tolA* deletion mutant so other strains were selected, as described above. The *tolRA* double deletion mutant JC8031 produced the highest amount of OMVs per cell, apart from *tolA* 838, so this strain was subsequently used for InaKtrBM3 production. Of note, from here on now, TB medium instead of LB medium was used when collecting OMVs of an exponential phase culture, as it was observed that higher cell densities were obtained in TB medium. A first expression test showed that InaKtrBM3 was produced and successfully surface displayed after a 3 h expression phase. Instead of performing a conversion assay, followed by GC-MS analysis, a quick and easy continuous spectrophotometric enzymatic assay was used for assessing activity. Schwaneberg et al developed an enzymatic assay, specifically for BM3 [31]. The assay is based on the conversion of *p*-nitrophenoxycarboxylic acids (pNCAs) by ω-hydroxylation, thereby releasing ω-oxycarboxylic acids and the chromophore p-nitrophenol. Assaying intact JC8031 cells after InaKtrBM3 production, the cells proved to produce p-nitrophenol (**Error! Reference source not found.**A). As a blank, NADPH was left out of the reaction mixture. NADPH is not able to traverse the membrane so observed activity upon NADPH addition must be the result of surface displayed InaKtrBM3. The OMVs were isolated and assayed as well, showing 12-pNCA conversion activity (**Error! Reference source not found.**B). An extra wash step was performed in order to remove any residual medium and these washed OMVs were again assayed (**Error! Reference source not found.**C). Unfortunately, no activity was observed anymore. The OMVs were further analysed by SDS-PAGE and compared to OMVs of JC8031, not producing any recombinant protein. In parallel, the OM fraction was isolated and taken along in the SDS-PAGE analysis (**Error! Reference source not found.**A). In the OM lanes, a band appears around 150 kDa, which is absent in case of JC8031, not producing recombinant protein. This band was not seen in the OMV lanes. The band was cut out and by MALDI-TOF MS it was confirmed to be InaKtrBM3 (data not shown). An expression test producing InaKtrBM3 overnight in LB medium, in analogy to the surface display experiments, was performed as well, showing similar results. Due to the fact that activity was lost upon extra washing of the OMV pellet, a medium sample was analysed by SDS-PAGE (**Error! Reference source not found.**B). Here, a band appeared at a slightly lower MW. Again, this band was cut out and analysed by MALDI-TOF MS. It was identified as BM3 and no InaKtr-specific peptides could be found in the MS spectrum (data not shown). Taken together with the aforementioned activity in the crude OMV sample, it was clear that a significant amount of active BM3 is cleaved of the InaKtr anchor and released in the medium.

During these experiments, it was evident that the surface display of InaKtrBM3 in JC8031 led to growth impairment as the optical density was significantly lower than the density reached by JC8031, not producing recombinant protein, in the same period of time. An OD_600_ of 2.15 was reached after a 3 h expression phase, compared to 5.16 for JC8031, not producing recombinant protein. After overnight expression, an OD_600_ of 2.72 was measured, compared to 6.12 for JC8031, not producing recombinant protein. This slower growth was already seen during the growth phase, indicating leaky expression. Efforts were done to resolve these issues. On the one hand, glucose was added and IPTG concentration for induction was reduced to 0.3 mM. On the other hand, *inaKtrBM3* was cloned into the vector pBAD/Myc-HisA, where the gene is under the control of the more tightly regulated arabinose-induced *araBAD* promotor. In both cases, optical densities were reached comparable to JC8031, with the construct under the control of the *araBAD* promotor showing superior growth over the IPTG-induced culture. Unfortunately, no OMVs were found to contain InaKtrBM3, neither after a 3 h expression phase, upon collection in the late exponential phase and after overnight expression (Figure 8A, B and C, respectively). From these experiments, it was confirmed that a significant amount of BM3 was cleaved of the InaKtr anchor and released in the medium. A concentrated sample of the OMV supernatant showed the most intense band at a lower MW than the expected MW of 152 kDa upon collection at late exponential phase (Figure 8B). Also after overnight expression, a lower MW band is observed in the supernatant (and in the crude OMV sample collected after expression of InaKtrBM3_pMAL) (Figure 8C).

**Figure 6:**
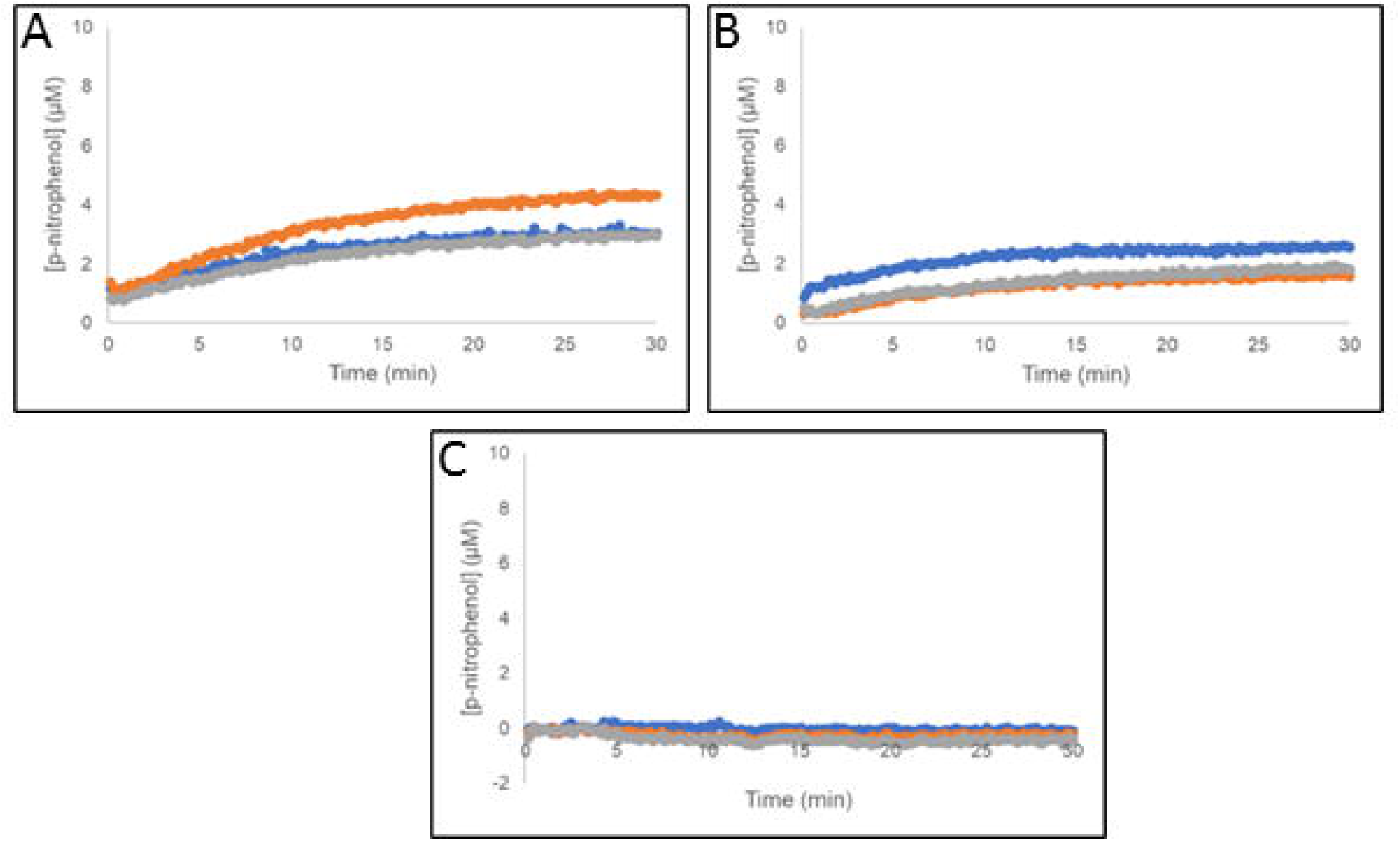
12-pNCA assay after InaKtrBM3 production in JC8031 for 3 h in TB medium. A: cells. B: crude OMVs. C: washed OMVs.

**Figure 7:**
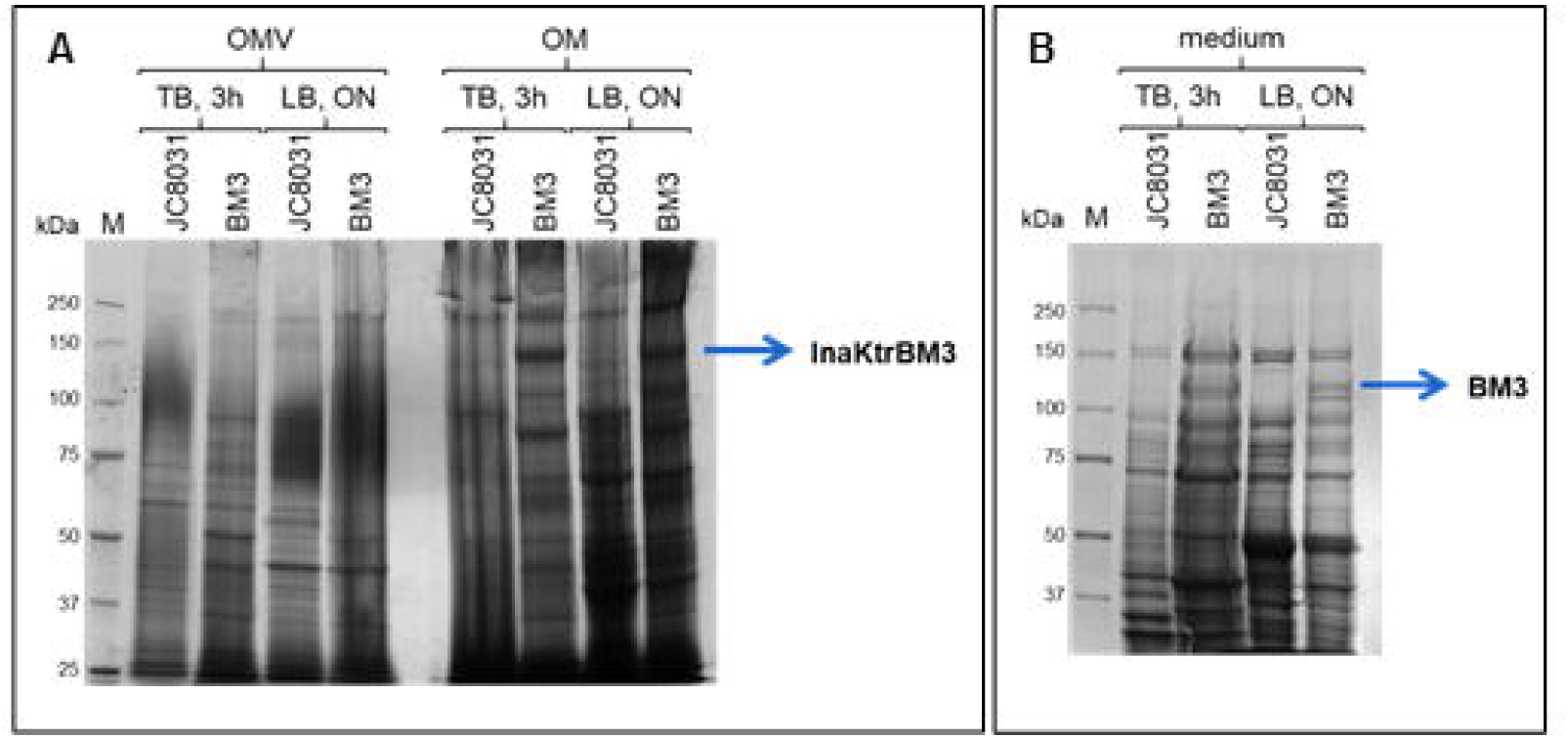
SDS-PAGE analysis after InaKtrBM3 production in JC8031 for 3 h in TB medium or overnight in LB medium. A JC8031 culture not producing recombinant protein was taken along in parallel. A: OMV and OM samples The band appearing around 150 kDa was identified as InaKtrBM3 by MALDI-TOF MS. B: Concentrated medium samples. The band appearing in between 100 kDa and 150 kDa was identified as BM3 and no peptides, specific for the InaKtr anchor were found.

**Figure 8:**
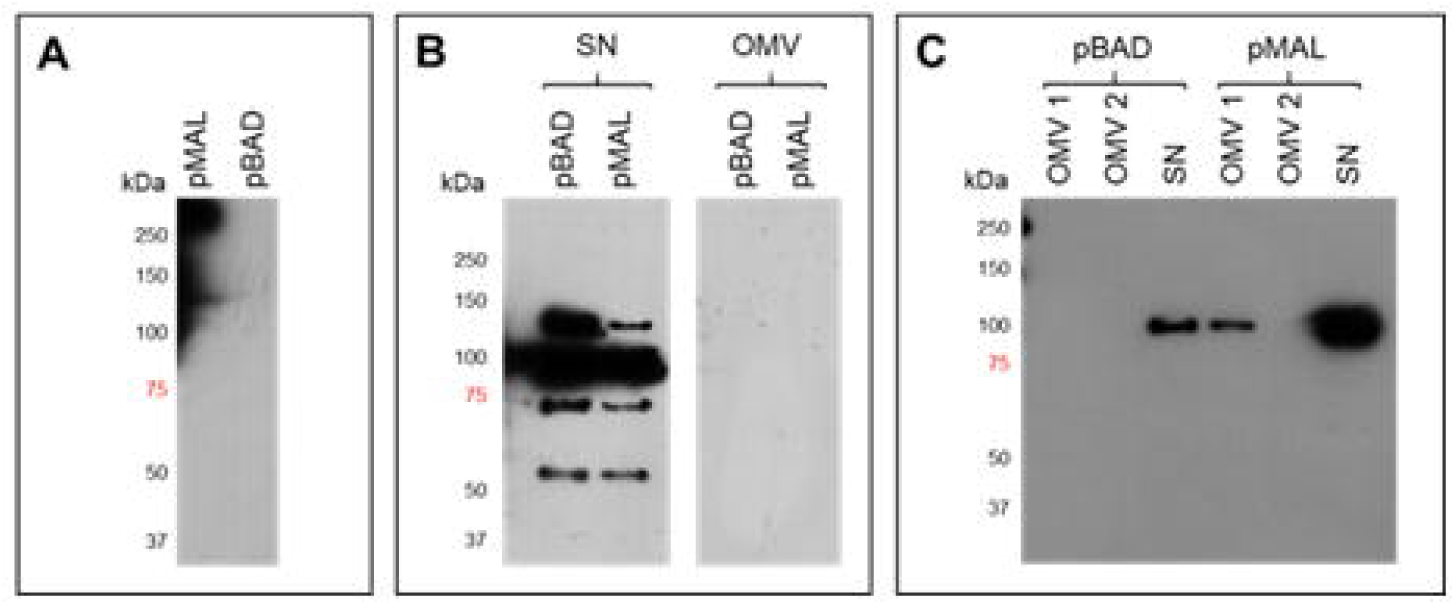
Western blot analysis of OMV samples derived from JC8031 after producting InaKtrBM3, either under the control of the *tac* promotor (pMAL) or under the control of the *araBAD* promotor (pBAD). A: 3 h expression phase. B: OMV collection in the late exponential phase. A concentrated sample of the OMV supernatant (SN) was taken along in parallel. C: Overnight expression phase. An OMV sample was taken before and after an extra wash step (i.e. OMV 1 and OMV 2, respectively), together with a concentrated sample of the OMV supernatant.

### 3.5 OMVs displaying InaKtrBM3 were obtained from JC8031 by EDTA-extraction

No spontaneously formed OMVs displaying InaKtrBM3 were isolated so far. Therefore, another path was taken, inspired by methods developed for application of OMVs as a vaccine. In the development of OMV-based vaccines, not only spontaneously formed OMVs are investigated (from here on now referred to as sOMV). Rather, OMVs are extracted after cell collection using either detergents, resulting in so-called dOMVs, or detergent-free extraction methods [15]. These detergent-free extraction methods are reported to result in more native-like OMVs (from here on now referred to as extracted OMVs or eOMVs). Van de Waterbeemd et al reported the use of EDTA, a chelating agent that destabilizes the OM and thereby releasing OMVs, in combination with a genetic modification resulting in a loosely attached OM. This approach resulted in a good eOMV yield, and this yield was superior to the sOMV yield [30]. Therefore, EDTA extraction of eOMVs from the JC8031 strain, recombinantly producing InaKtrBM3, was performed. Both the pMAL and the pBAD constructs were taken along in parallel, as well as the original JM109 strain, surface displaying InaKtrBM3. After isolation of the eOMVs by ultracentrifugation, followed by a wash step, the eOMVs were assayed for activity (Figure 9). Expression of *inaKtrBM3* under the control of the *tac* promotor (pMAL) led to eOMVs showing p-nitrophenol formation to equilibration (Figure 9A). A moderate activity was seen in case of arabinose induction (pBAD) (Figure 9B). No activity was seen when producing InaKtrBM3 in the JM109 strain (Figure 9C). Indeed, western blot analysis shows that InaKtrBM3 is present in both JC8031-derived eOMVs, whereas the enzyme is absent from eOMVs, derived from JM109 (Figure 9D).

**Figure 9:**
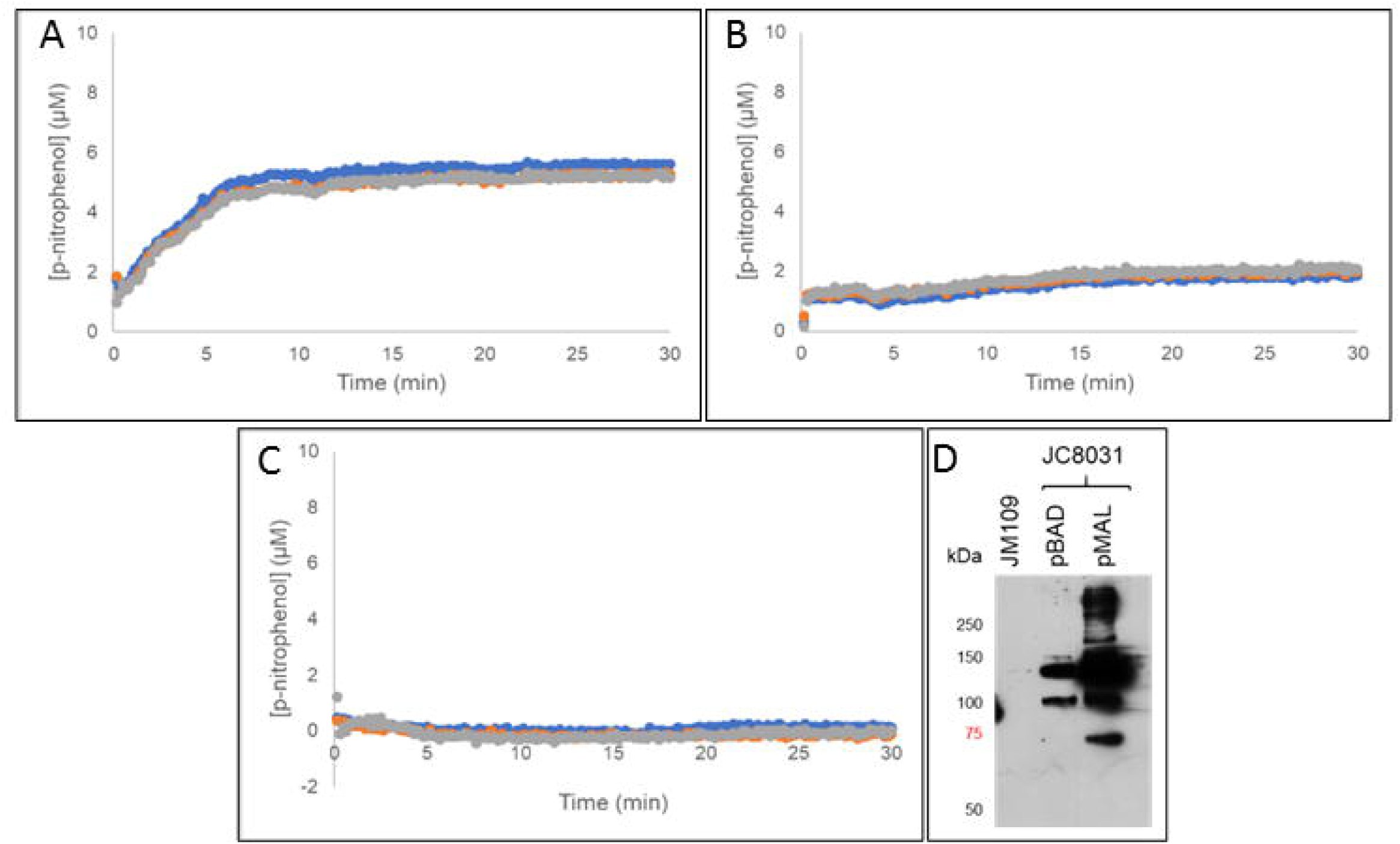
eOMVs extracted after InaKtrBM3 production. A: 12-pNCA assay. JC8031-derived eOMVs. Expression was under the control of the *tac* promotor (pMAL). B: 12-pNCA assay. JC8031-derived eOMVs. Expression was under the control of the *araBAD* promotor (pBAD). C: 12-pNCA assay. JM109-derived eOMVs. D: Western blot analysis.

### 3.6 eOMVs, but not sOMVs, displaying InaKtrBM3 were isolated from both JC8031 and the *nlpI* deletion mutant

Although success was booked with eOMVs, interest still went to obtaining sOMVs, as collection of these vesicles does not require an extra extraction step. To this end, both InaKtrBM3_pMAL and InaKtrBM3_pBAD were transformed in the *nlpI* deletion mutant (*nlpI* 863). After overnight expression, allowing the cells to grow to the stationary phase (as only then, hypervesiculation was observed), both sOMVs and eOMVs were collected. sOMVs were collected from the JM109 strain as well, where hypervesiculation was induced by polymyxin B addition. eOMV extraction from the JM109 strain was again performed, but now after overnight expression. The JC8031 strain was taken along in parallel. In case of JC8031, the eOMVs were clearly active (**Error! Reference source not found.**A and B), with the eOMVs after IPTG induction reaching equilibrium very quickly (**Error! Reference source not found.**A). The sOMVs derived from JC8031 seemed to show minor activity (**Error! Reference source not found.**C and D). OMVs derived from the JM109 strain did not show any activity (**Error! Reference source not found.**B). However, some minor activity was seen in the eOMV sample (**Error! Reference source not found.**A). InaKtrBM3 was present in the eOMVs derived from *nlpI* 863 after IPTG induction, as a small amount of p-nitrophenol was formed (Figure 12A). Also the eOMVs after arabinose induction seem to be slightly active (Figure 12B). Neither of the sOMV samples showed any p-nitrophenol formation. These experiments show that the extraction of OMVs using EDTA, in combination with mutations impairing envelope stability, result in the highest OMV yields.

**Figure 10:**
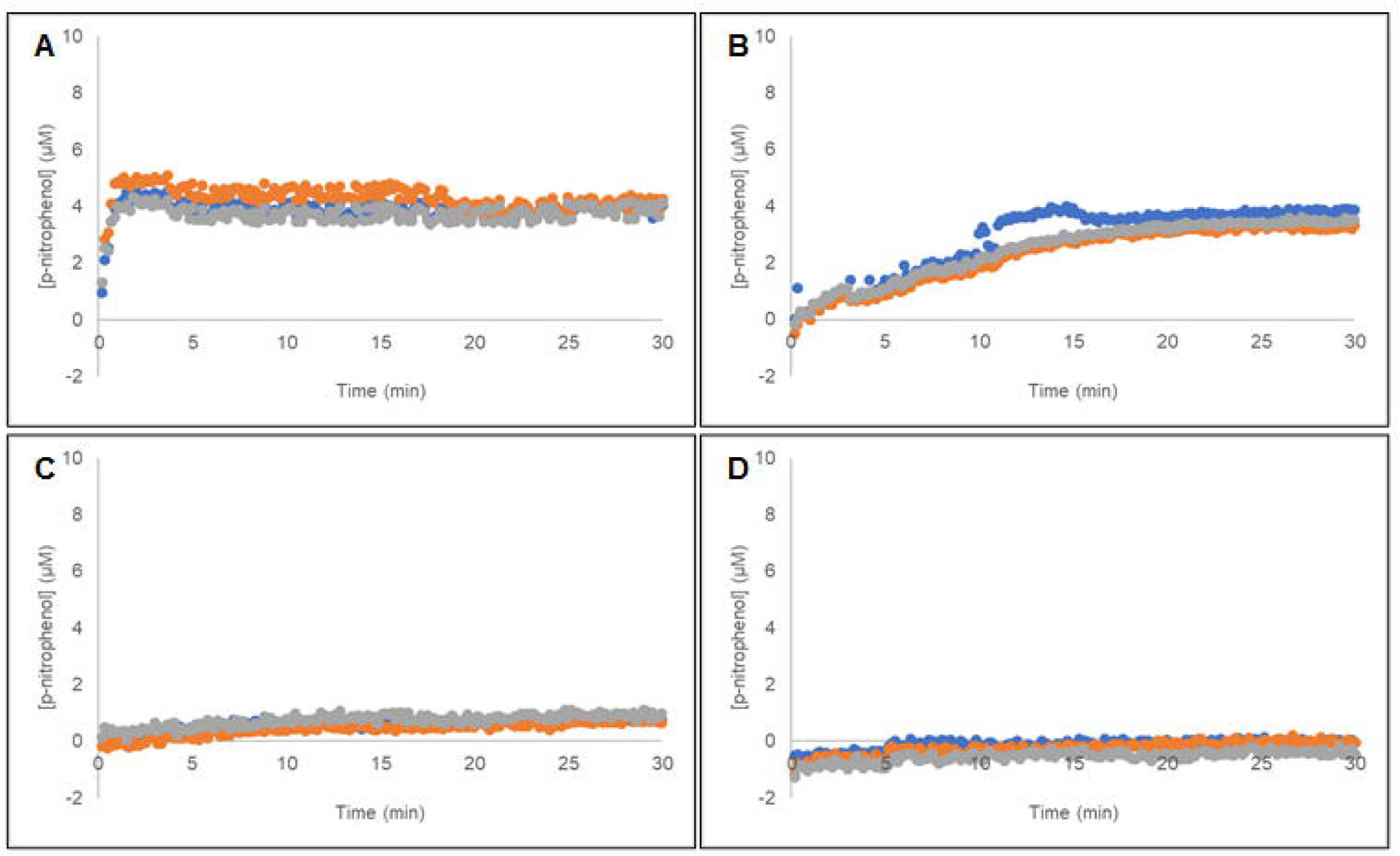
12-pNCA assay using OMVs isolated after InaKtrBM3 production in JC8031. A: eOMVs. Expression was under the control of the *tac* promotor. B: eOMVs. Expression was under the control of the *araBAD* promotor. C: sOMVs. Expression was under the control of the *tac* promotor. D: sOMVs. Expression was under the control of the *araBAD* promotor.

**Figure 11:**
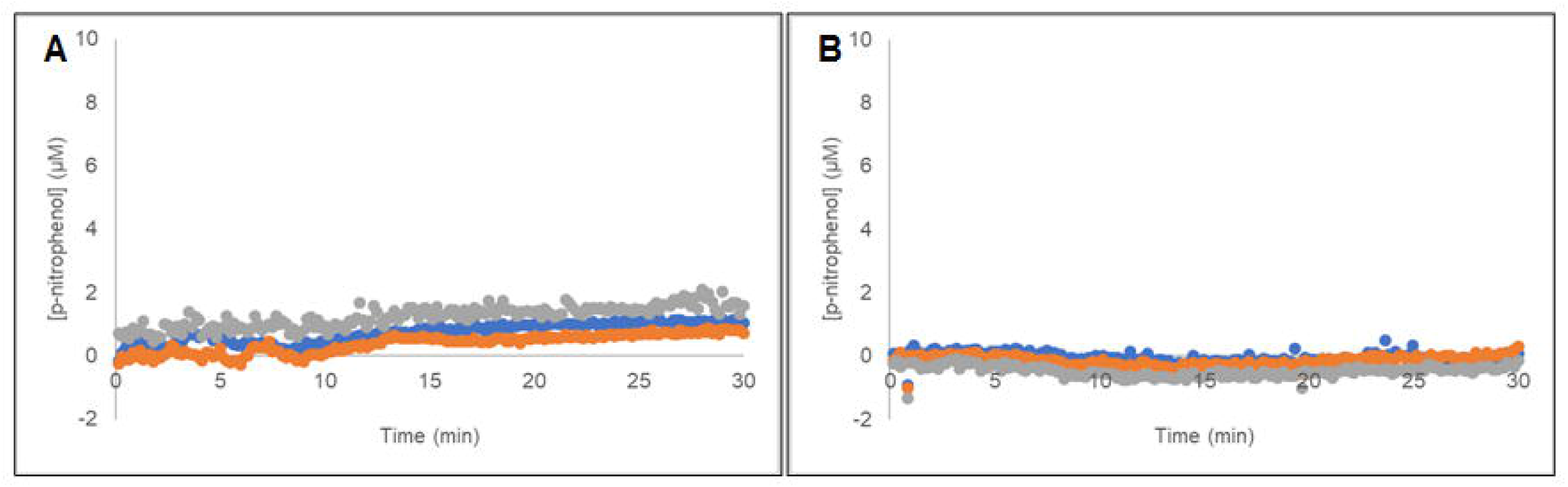
12-pNCA assay using OMVs isolated after InaKtrBM3 production in JM109. A: eOMVs extracted after overnight expression. B: sOMVs collected 6 h post polymyxin B induction.

**Figure 12:**
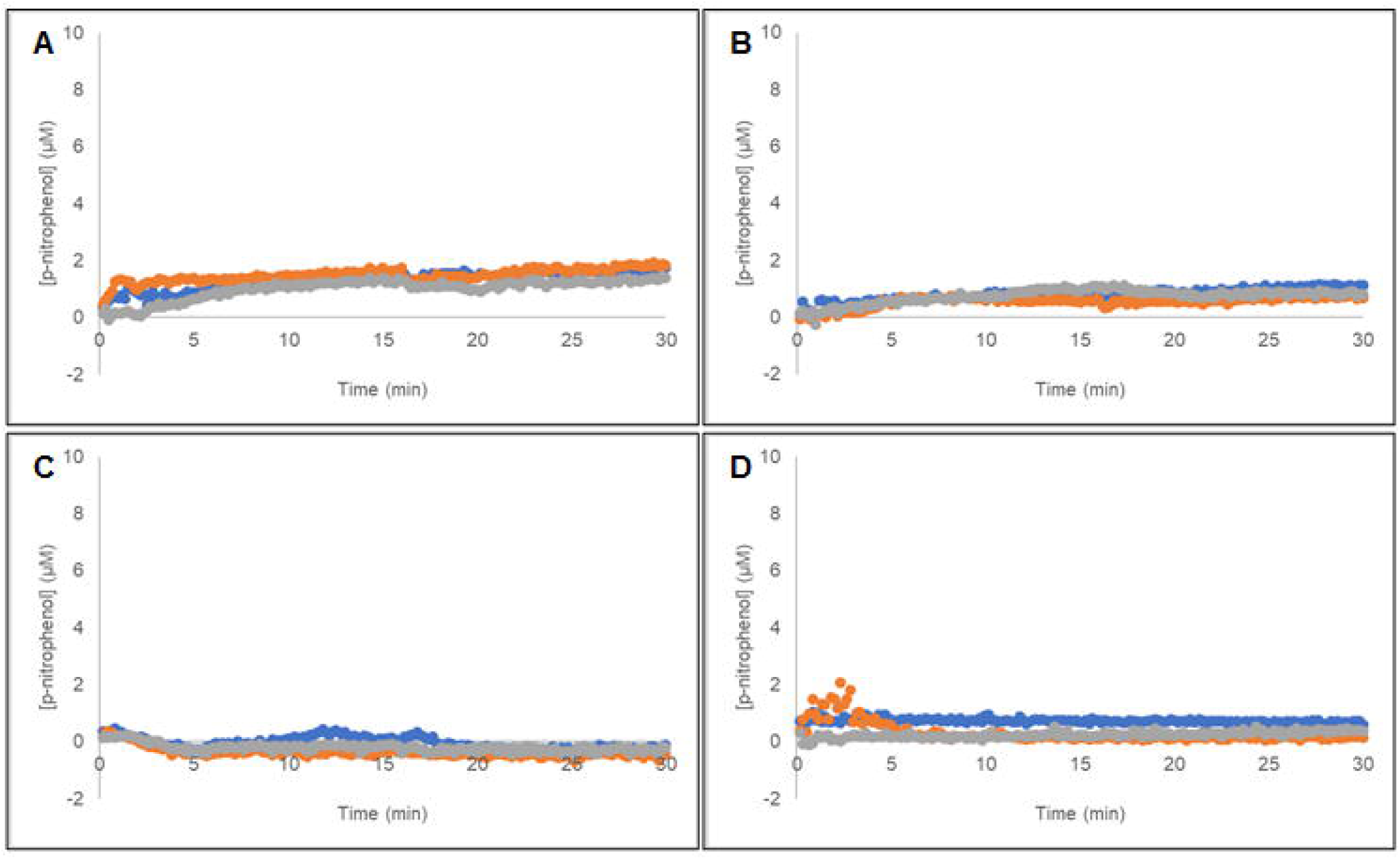
12-pNCA assay using OMVs isolated after InaKtrBM3 production in *nlpI* deletion mutant 863. A: eOMVs. Expression was under the control of the *tac* promotor. B: eOMVs. Expression was under the control of the *araBAD* promotor. C: sOMVs. Expression was under the control of the *tac* promotor. D: sOMVs. Expression was under the control of the *araBAD* promotor.

### 3.7 sOMVs displaying InaKtrBM3 could be isolated from JC8031 when exchanging the 0.22 µm filter for a 0.45 µm filter

So far, the culture supernatant was filtered using a 0.22 µm filter before sOMV collection by ultracentrifugation, as this was an established protocol within the lab. This pore size is smaller than the upper limit of the OMV size. However, scanning literature learned that 0.45 µm filters are often used instead of 0.22 µm filters [18], [36]. We wondered whether the use of 0.45 µm filters could increase the OMV yield, and thereby obtaining a sufficient amount of active OMVs. A culture was set up where InaKtrBM3 was produced in JC8031 upon IPTG induction. The culture supernatant was divided in two. One half was filtered using a 0.22 µm filter, whereas the other half was filtered using a 0.45 µm filter. In this way, it was made sure that no differences could be ascribed to biological variations between different cultures. The isolated sOMVs were subsequently assayed for 12-pNCA hydroxylation. An increasing absorbance was observed after use of the 0.45 µm filter, whereas no such increase was seen in case of the 0.22 µm filter.

As InaKtrBM3 seemed to be present in sOMVs, collected after filtering the supernatant with a 0.45 µm filter, all strains and constructs, mentioned above, were reinvestigated. Both eOMVs and sOMVs were collected, now filtering the respective supernatant with a 0.45 µm filter, and tested for activity by the 12-pNCA assay. First, the JC8031 strain was revisited in order to confirm that the sOMVs, obtained after 0.45 µm filtration indeed were active and the arabinose-induced construct was taken along in parallel (Figure 14). In Figure 14A, p-nitrophenol formation was clearly seen, thereby confirming that active sOMVs were successfully obtained. It must be noted that the medium appeared to be somewhat viscous and eOMV extraction was no longer possible as cells lysed upon incubation with the EDTA-rich buffer. It was noticed that every time the JC8031 strain with the construct under the control of the *tac* promotor was grown, starting from a glycerol stock made from the preceding preculture, a slightly lower OD_600_ value was reached upon collection in late exponential phase. This raised the question whether these OMVs truly are sOMVs resulting from OM blebbing, or whether these are the result of (partial) cell lysis. There was no clear activity seen for any other sOMV sample (Figure 14B and D). The JC8031-derived eOMVs collected after InaKtrBM3 production, under the control of *araBAD* were active (Figure 14B), as well as the *nlpI* 863-derived eOMVs collected after InaKtrBM3 production under the control of the *tac* promotor (Figure 14A), which was also seen after filtration with a pore size of 0.22 µm. Now, also the *nlpI* 863-derived eOMVs under the control of *araBAD* showed activity (Figure 14C). Neither the vesicles derived from *tolA* 838 nor derived from the polymyxin B-induced JM109 strain actively converted 12-pNCA (data not shown).

**Figure 13:**
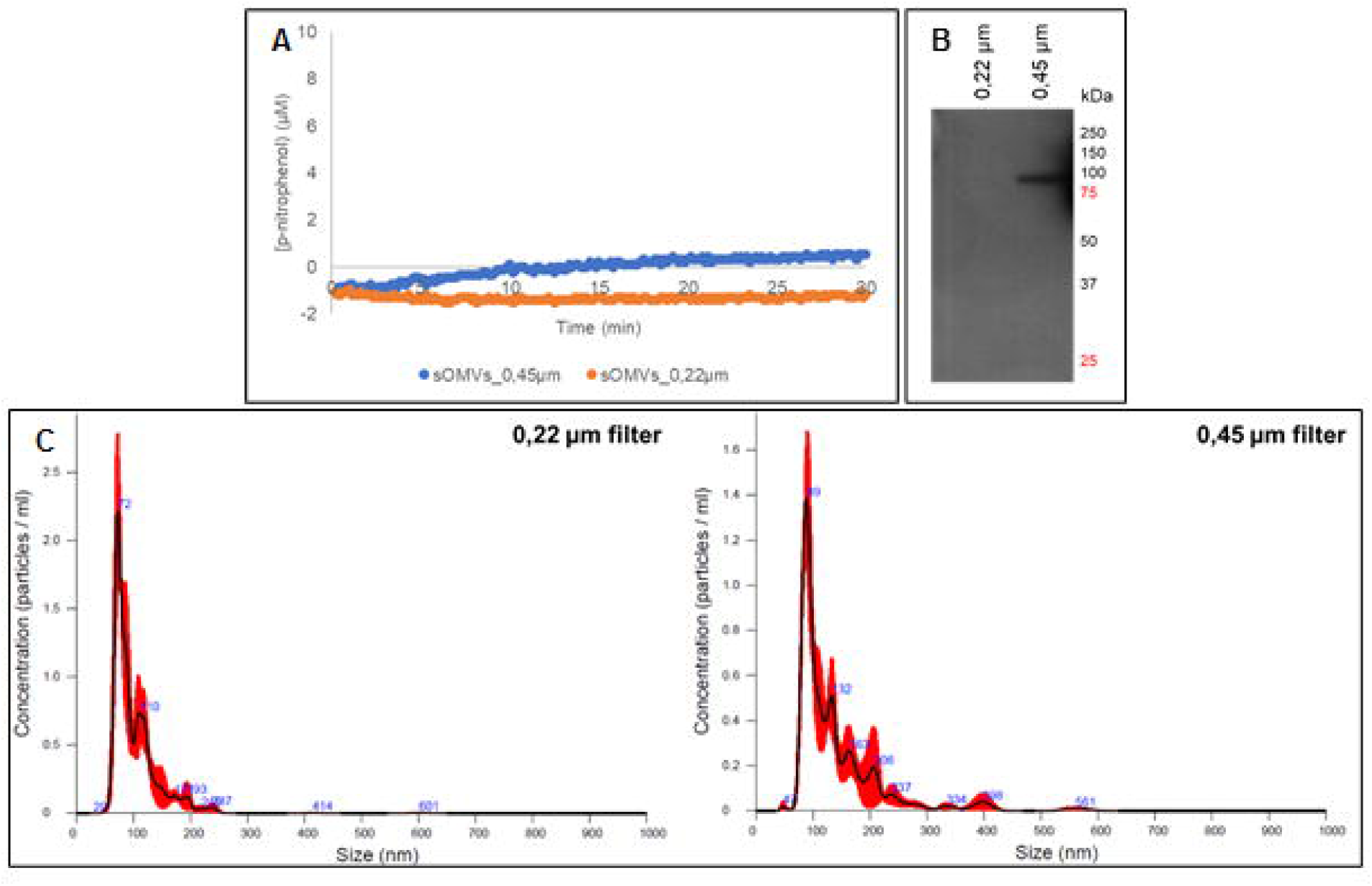
sOMVs isolated after InaKtrBM3 expression in JC8031 under the control of the *tac* promotor. One half of the medium was filtered with a 0.22 µm filter. The other half was filtered with 0.45 µm filter. A: 12-pNCA assay. B: Western blot analysis. C: NTA analysis.

**Figure 14:**
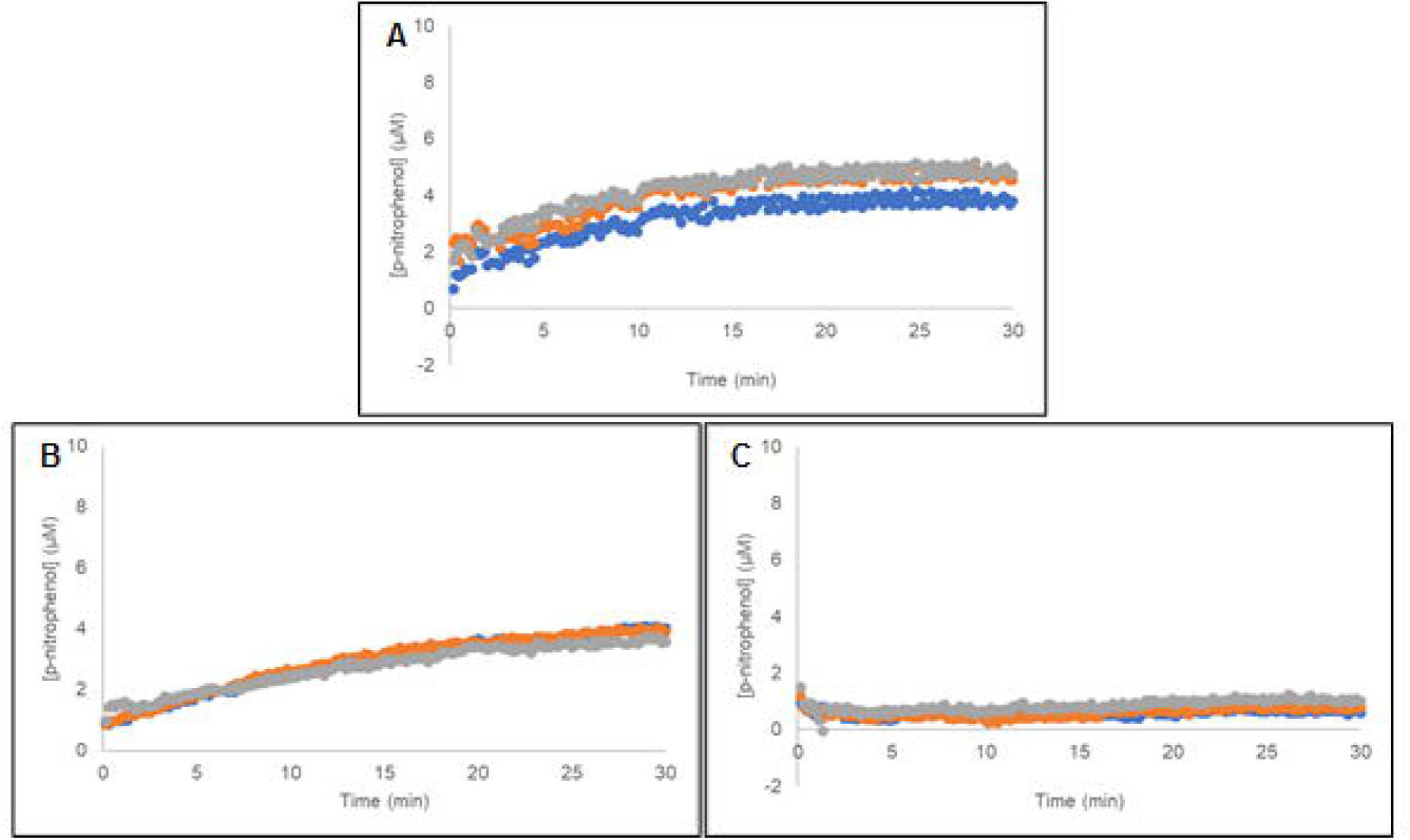
12-pNCA assay using OMVs isolated after InaKtrBM3 expression in JC8031. Filtration was done with a 0.45 µm filter instead of a 0.22 µm filter. A: sOMVs. Expression was under the control of the *tac* promotor. B: eOMVs. Expression was under the control of the *araBAD* promotor. C:sOMVs. Expression was under the control of the *araBAD* promotor.

**Figure 15:**
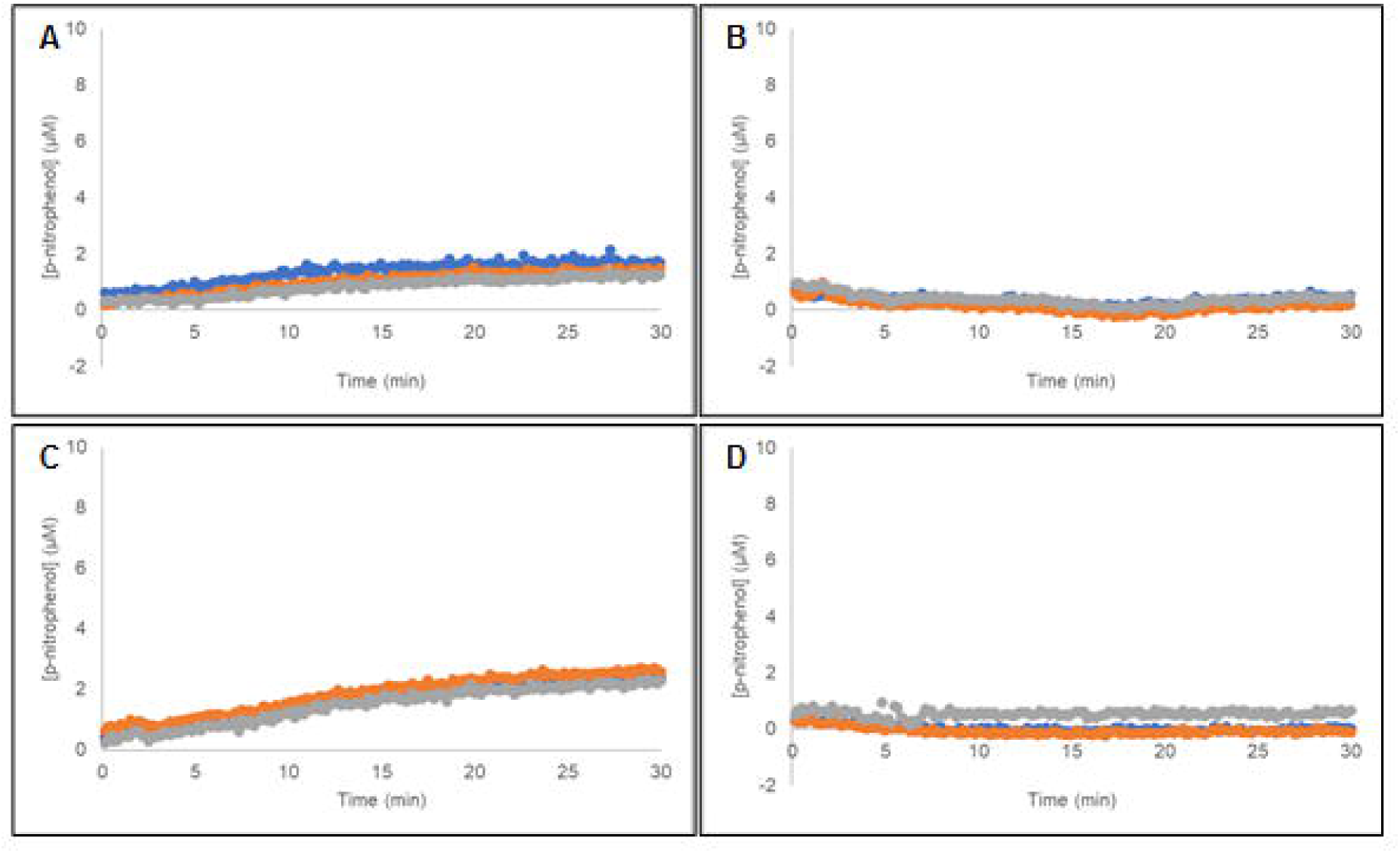
12-pNCA assay using OMVs isolated after InaKtrBM3 production in *<u>nlpI</u>* deletion mutant 863. Filtration was done with a 0.45 µm filter instead of a 0.22 µm filter. A: eOMVs. Expression was under the control of the *tac* promotor. B: sOMVs. Expression was under the control of the *tac* promotor. C: eOMVs. Expression was under the control of the *araBAD* promotor. D: sOMVs. Expression was under the control of the *araBAD* promotor.

## 4 Discussion

An interesting approach to combine advantages of both whole-cell bioconversions and *in vitro* biocatalysis, is the surface display of CYP enzymes on the outer membrane (OM) of *E. coli*. On the one hand, stability of the produced CYP is enhanced by *in vivo* immobilization into a membrane environment and no expensive enzyme purification process is required. On the other hand, no membrane barrier needs to be crossed, possibly leading to higher productivities [37]. In this article, it was investigated if this surface display approach could be taken one step further, i.e. displaying a self-sufficient CYP on OMVs derived from the *E. coli* OM. Thereby, a true *in vitro* cell-free system is created, eliminating possible unwanted side reactions occurring in the cellular environment. Park et al showed how OMV display led to an increased activity compared to yeast surface display of the same enzyme complex [16]. It was hypothesized that the increased activity could be ascribed to the nanoscale dimensions of the OMVs, resulting in an increased enzyme:volume ratio and improved substrate accessibility [16], [17]. In this article, membrane vesicles were successfully isolated, either EDTA-extracted eOMVs or spontaneously formed sOMVs and showed to display CYP102A1, actively converting the model substrate 12-pNCA. OMVs have proven their value in the development of a vaccine against *Neisseria meningitis* and are increasingly investigated as a vaccine against other pathogens [15]. Furthermore, the use of engineered OMVs has emerged as vehicles for targeted drug delivery [11]. In this article, it was shown that OMVs can serve as an *in vivo* immobilization strategy for recombinant production of self-sufficient CYP monooxygenases, thereby proposing an alternative production strategy.

OMVs displaying an active self-sufficient CYP enzyme were successfully isolated. EDTA-extracted eOMVs were obtained in four different cases, i.e. using one of two deletion mutants JC8031 or *nlpI* 863 in combination with either the pMAL or pBAD vector. This confirms the study in which Van der Waterbeemd showed that OMV extraction with EDTA from a gram-negative bacterium, genetically modified resulting in a loosely attached OM, yields a higher amount of vesicles compared to the isolation of spontaneously formed sOMVs [30]. Spontaneously formed sOMVs were isolated only when the *tolRA* double deletion mutant JC8031 was used to express the InaKtrBM3-encoding gene under the control of the *tac* promotor in the pMAL vector. It became evident that obtaining sOMVs in a sufficient yield to observe activity, requires a very delicate equilibrium between OM instability and viability. In the first experiments, the strain transformed with the pMAL construct showed good optical densities at the point of OMV collection. Unfortunately, every consecutive round of OMV production, starting from a glycerol stock made during the previous experiment, a decreased optical density was reached. At a certain point, eOMVs could no longer be extracted as the cells lysed upon contact with the EDTA-buffer. This exemplifies this delicate equilibrium. A construct under the control of the *araBAD* promotor did not result in a sOMV yield, sufficient to observe activity. This promotor is proposed to be used in order to better control expression, however this negatively affects the sOMV yield. Knocking out *nlpI* was reported to result in hypervesiculation without affecting the membrane integrity. Indeed hypervesiculation was observed, but only after overnight growth. However, this increased membrane integrity negatively affected the sOMV yield. It thus seems that the membrane instability really needs to be pushed to its limits and a strong promotor is required for sOMV production. One might therefore wonder if these are indeed all true sOMVs or if cell lysis occurs and results in released membrane fractions containing the surface displayed construct. Indeed, it must be noted that knock-out strains with disrupted cross-links, often showed a leaky membrane and Toyofuku et al reported that vesicles formed by cell lysis might be co-isolated in that case [19]. Of note, the *tolA* deletion mutant showed a similar hypervesiculating phenotype as JC8031, but did not result in OMVs containing InaKtrBM3, although the enzyme was present in the OM fraction. It must be considered that the isolated OM fraction might rather be an OM enrichment and might therefore still contain enzymes from other cellular fractions.

Throughout this work, an enzymatic assay with the substrate 12-pNCA was used in order to measure the activity spectrophotometrically in case of InaKtrBM3 display. This proved to be a valuable alternative to GC-MS. However, it was observed that only a fraction of the added substrate seemed to be converted to p-nitrophenol. It must be noted that only hydroxylation of 12-pNCA at the ω-position will result in the chromophore p-nitrophenol. BM3 preferentially hydroxylates the subterminal positions ω-1, ω-2 and ω-3. Products resulting from hydroxylation at a position other than the terminal position will not be visible by spectrophotometry, explaining the low yield of p-nitrophenol. Using the F87A mutant, the conversion of 12-pNCA to p-nitrophenol was much higher, as reported by Schwaneberg et al, so this this mutant might be used instead to enhance sensitivity [31].

The OMV display approach shows great promise, although there are still many parameters that need to be investigated and optimized before this would result in an economically attractive process.

Firstly, optimization of the linker sequence fusing the anchor to the enzyme should be addressed in further application. The linker sequence was sufficient to retain enough enzyme at the surface of either the cell or the OMV in order to obtain an actively converting immobilized CYP. Unfortunately, a high amount of BM3 enzyme was found in the culture medium, indicating that the linker sequence is susceptible to proteolysis. Although the released BM3 was still active, this has implications for the costs of the catalytic process. Using either cells or OMVs gives the opportunity for easy catalyst recycling. However, when the enzyme is cleaved of the anchor, the activity of the recovered catalyst will significantly reduce at each recycling step.

Secondly, it is of interest to obtain actual sOMVs, obtained from budding of the OM, as this greatly facilitates downstream processing. In this context, other hypervesiculating mutants might be investigated as well. For example, not only reduced crosslinking, but also envelope stress was reported to be involved in OMV biogenesis. Disruption of genes involved in the σ^ E^ stress response pathway, that is *degS*, *degP* and *rseA*, led to a marked OMV increase. This pathway is activated in response to misfolded protein in the periplasm and it was hypothesized that OMV formation was a result of protein accumulation, leading to bulging of the OM and finally secreting these accumulated proteins through OMVs [38]. Of note, next to InaK, other INPs have been identified and it was reported that alternative INPs differ in their efficiency to target their recombinant protein passenger to the surface [26]. Also Autodisplay resulted in successful display of CYPs on the surface of *E. coli* [10, 39]. Increasing the yields of active sOMVs should therefore also include investigation of alternative OM anchors, next to alternative hypervesiculating strains.

Thirdly, the stability of these OMVs needs to be assessed. OMVs have been reported to be stable during long-term storage and upon several cycles of freeze-thawing [40], [41]. However, the stability in the desired biocatalytic setup where e.g. shearing forces and solvent will be present, needs to be tested.

Fourthly, it might be of interest to engineer *E. coli* for the inclusion of a heme biosynthetic pathway. The heme precursor δ-aminolevulinic acid (5-ALA) is expensive so elimination of this additive would greatly reduce the production costs. For example, Harnastai et al showed that co-expression of glutamyl-tRNA reductase proved to be sufficient for eliminating the need for 5-ALA supplementation [43].

Another parameter that needs attention is the purification of these vesicles. OMVs are now generally collected by ultracentrifugation or ultrafiltration techniques. Alternatively, it has been shown that IMAC can be used for the purification of OMVs [44], thereby only retaining vesicles displaying the protein of interest. On an industrial scale, both ultrafiltration and IMAC are established but expensive purification processes.

Therefore, it remains to be verified if OMVs would be a suitable nanobiocatalyst. For example in the pharmaceutical industry, CYPs are involved in the production of many interesting compounds. Moreover, liver CYP enzymes are of major importance in drug metabolism and great interest continues to go to recombinant production systems as biosensors in drug development. OMV display is thus believed to serve as an invaluable immobilization strategy for these applications.

The cofactor NADPH is required for hydroxylation and to make a biocatalytic process economically feasible, a cofactor regeneration system should be included. The OMV display approach offers the possibility to create a true self-sufficient biocatalyst, by co-expression and OMV display of both the self-sufficient CYP and a regeneration system (e.g. formate dehydrogenase, glucose dehydrogenase, alcohol dehydrogenase or phosphite dehydrogenase). Furthermore, co-expression would allow for an approach where all enzymes are isolated in one and the same purification process. A fusion construct between CYP102A1 and PTDH was created as well in the search for a true self-sufficient enzyme [45].

## 5 Conclusion

Immobilization is a well-known strategy to increase stability of a biocatalyst. In order to circumvent a costly purification before continuing to immobilization, an *in vivo* immobilization strategy was proposed, targeting self-sufficient CYPs to the surface of OMVs. A proof-of-concept was delivered using the model enzyme CYP102A1, by either isolating active sOMVs from the medium or extracting active eOMVs from the cells. The origin of the sOMVs, i.e. are the vesicles indeed spontaneously formed by blebbing of the OM or are these the result of cell lysis, requires further investigation, as well as their feasibility as a vehicle for large scale application. However, it is believed that OMV display shows great promise in other applications, e.g. in the development of biosensors.

## Acknowledgments

This work was supported by a Catalisti-ICON project with funding from the Flemish Government via Vlaams Agentschap Innovatie en Ondernemen (VLAIO) to DD and BD. We thank Laetitia Houot and Inge Van Bogaert for providing strains or plasmids as described in Materials and Methods.

## Notes

### Competing Interest Statement

The authors have declared no competing interest.

