## Supplementary material for "Display of the self-sufficient CYP102A1 on the surface of *E. coli*-derived Outer Membrane Vesicles": Fig. S1

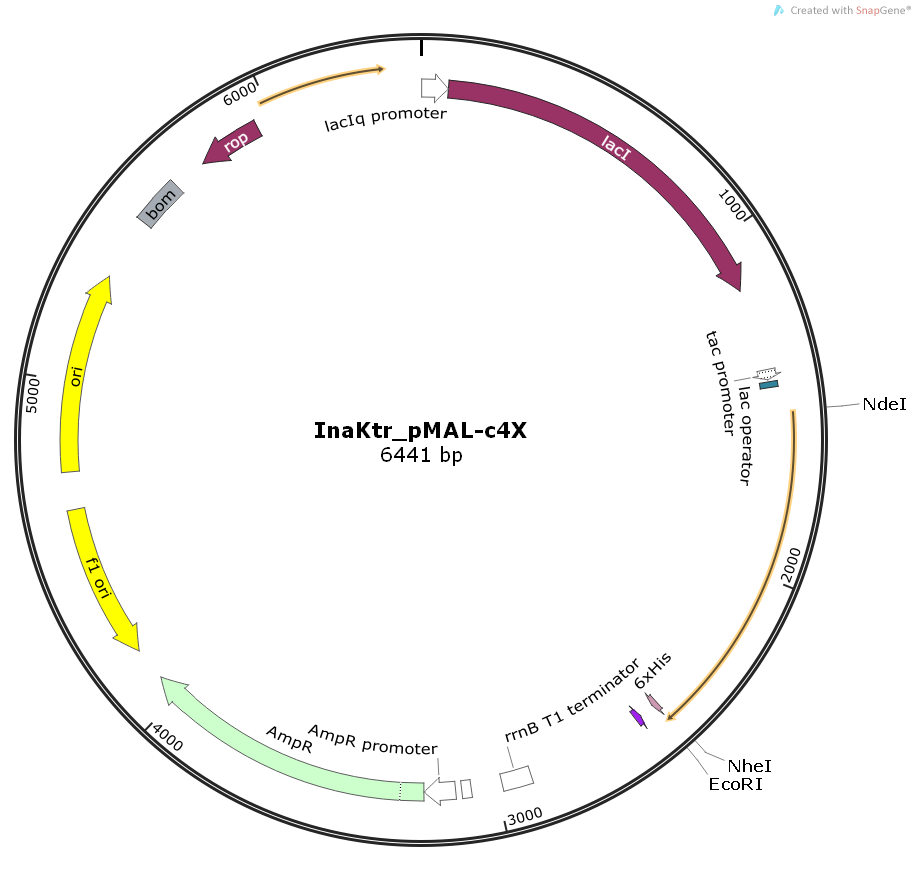

Supplementary Figure. Structure of the plasmid construct used for IPTG induced expression of the His tagged InaK-BM3 fusion.
